# Genetic variation in *Loudetia simplex* supports the presence of ancient grasslands in Madagascar

**DOI:** 10.1101/2023.04.07.536094

**Authors:** George P. Tiley, Andrew A. Crowl, Tchana O. M. Almary, W. R. Quentin Luke, Cédrique L. Solofondranohatra, Guillaume Besnard, Caroline E.R. Lehmann, Anne D. Yoder, Maria S. Vorontsova

## Abstract

Research Aims — The extent of Madagascar’s grasslands prior to human colonization is unresolved. We used population genetic analyses of a broadly dominant C_4_ fire-adapted grass, *Loudetia simplex*, as a proxy for estimating grassland change through time. We carefully examined the utility of target-enrichment data for population genetics to make recommendations for conservation genetics. We explored the potential of estimating individual ploidy levels from target-enrichment data and how assumptions about ploidy could affect analyses.
Methods — We developed a novel bioinformatic pipeline to estimate ploidy and genotypes from target-enrichment data. We estimated standard population genetic summary statistics in addition to species trees and population structure. Extended Bayesian skyline plots provided estimates of population size through time for empirical and simulated data.
Key Result — All Malagasy *Loudetia simplex* individuals sampled in this study formed a clade and possibly indicated an ancestral Central Highland distribution of 800m in altitude and above. Demographic models suggested grassland expansions occurred prior to the Last Interglacial Period and supported extensive grasslands prior to human colonization. Though there are limitations to target-enrichment data for population genetic studies, we find that analyses of population structure are reliable.
Key Point —Genetic variation in *Loudetia simplex* supports widespread grasslands in Madagascar prior to the more recent periods of notable paleoclimatic change. However, the methods explored here could not differentiate between paleoclimatic change near the Last Glacial Maximum and anthropogenic effects. Target-enrichment data can be a valuable tool for analyses of population structure in the absence a reference genome.

**Societal Impact Statement:** Recognizing *Loudetia* dominated grasslands were widespread prior to human colonization highlights that open ecosystems were and continue to be an important component of Madagascar’s biodiversity. Urgently required are biodiversity inventories and integrative taxonomic treatments of grassland flora and fauna to asses risks to understudied ecosystems historically regarded as wastelands. Substantial financial and logistical barriers exist to implementing conservation studies using contemporary genomic tools. We ameliorated some of the challenges for population genetic analyses of non-model polyploids lacking reference genomes by developing computational resources to leverage a cost-effective data generation strategy that requires no prior genetic knowledge of the target species.

**Résumé:**

1. Les objectifs de la recherche — L’étendue des écosystèmes ouverts de Madagascar avant la colonisation humaine reste à éclaircir. Nous avons utilisé une analyse de la population génétique d’une graminée C_4_ adaptée au feu, largement dominante, Loudetia simplex, comme référence pour estimer les changements au niveau de ces biomes au fil du temps. Nous avons examiné attentivement l’utilité des données d’enrichissement ciblé pour la génétique de population afin de formuler des recommandations pour la conservation génétique. Nous avons exploré le potentiel de l’estimation du niveau des ploidies individuelles à partir des données d’enrichissement ciblé et comment les hypothèses à propos de ces ploidies pourraient affecter les analyses.
2. Les méthodes — Nous avons développé un nouveau canal bioinformatique pour estimer les ploidies et les génotypes à partir des données d’enrichissement ciblé. Nous avons estimé les statistiques standard de la population génétique, en plus des arbres des espèces et de la structure de la population. L’utilisation des tracés étendus du ciel bayésien a fourni une estimation de la taille de la population au fil du temps pour des données empiriques et simulées.
3. Résultat clé — Tous les individus Malagasy de *Loudetia simplex* échantillonnés dans cette étude ont formé un clade, indiquant une éventuelle ancienne distribution dans les hauts plateaux. Les modèles démographiques suggèrent une expansion des prairies bien avant la dernière période interglaciaire et soutiennent l’existence d’une vaste distribution avant la colonisation humaine. Bien qu’il y ait des limites à l’enrichissement des données cibles pour l’étude de la génétique des populations, nous constatons que l’analyse des structures des populations est fiable.
4. Les points clés — La variation génétique de *Loudetia simplex* soutient l’existence de vastes prairies à Madagascar avant les périodes plus récentes de changements paléoclimatiques notables. Cependant, les méthodes explorées ici n’ont pas permis de faire la différence entre les changements paléoclimatiques près du dernier maximum glaciaire et les effets anthropogènes. Les données d’enrichissement ciblé peuvent être un outil précieux pour les analyses de la structure des populations en l’absence d’un génome de référence.

**Déclaration d’impact societal:** Reconnaître que les prairies dominées par Loudetia étaient répandues avant la colonisation humaine souligne que les écosystèmes ouverts étaient et continuent d’être un composant important de la biodiversité de Madagascar. Il est urgent de réaliser des inventaires de la biodiversité et une taxonomie intégrée pour le traitement de la flore et de la faune des écosystèmes ouverts afin d’évaluer les risques pour les écosystèmes sous-étudiés considérés historiquement comme des terres en friches. Des barrières financières et logistiques existent pour mettre en œuvre l’étude de la conservation en utilisant les outils génomiques contemporains. Nous avons amélioré certains des défis liés aux analyses génétiques de populations de polyploïdes non modèles, sans génomes de référence, en développant des ressources informatiques pour exploiter une stratégie pouvant générer des données rentables ne nécessitant aucune connaissance génétique préalable de l’espèce cible.

**Famintinana:**

1. Ny tanjon’ny fikarohana — Mbola tsy fantatra mazava tsara ny fivelaran’ny hivoka teto Madagasikara talohan’ny fahatongava’ny olombelona. Mba ahafantarana ny fihovana nitranga nandritra ny fotoana naharitra teo amin’ireo hivoka ireo dia nanao famakafakahana ara-genetika amin’ny ahitra C_4_ miompana amin’ny afo iray antsoina Loudetia simplex ara-tsiantifika na Berambo na Hara amin’ny teny malagasy izahay. Nandinika tsara ny maha-zava-dehibe ny fampitomboana ny antotan-kevitra mba ahafahana manolo-kevitra momba ny fiarovana ny fototarazo genetika. Nandinika ny mety mampiavaka ny fanombanana an’ny ploidy tsirairay amin’ny fampitomboana antotan-kevitra sy ny mety ho fiantraikan’ny fiheverana momba ireo ploidy ireo amin’ny fikarohana.
2. Fomba Fiasa — Namorona fantsona bioinformatika vaovao mba ahafahana manombana ny ploidy sy ny « genotypes » avy amin’ny antotan-kevitra nokendrena izahay. Notombanana ny antontan’isa famintinana ny fototarazo ara-genetikan’ireo vondron’ahitra ireo, miampy ny karazana hazo sy ny firafitry ny vondrona na koa hoe mponina. Nanome tombantombana ny haben’ny mponina amin’ny alàlan’ny fotoana ny antontan-kevitra voavinavina azo tamin’ny fikarohana. Fikarohana izay azo tamin’ny alalan’ny « Bayesina Skuline Plots ».
3. Vokam-pikarohana fototra — Ny vondrona *Loudetia simplex* eto Madagasikara izay niasana dia namorona « clade » na fikambanana iray, izay manondro ny mety maha ela netezana sy tranainy an’io ahitra io eny amin’ny faritra avo. Ny modely demografika dia manoro hevitra amin’ny naha be velarana ny hivoka izay efa ela talohan’ny vanim-potoana « interglacial » farany ary manohana ny fivelarana midadasika an’ireo kijana ireo alohan’ny fonenan’ny olombelona. Na dia misy fetrany aza ny fampitomboana ny antotan-kevitra kendrena amin’ny fandalinana ny fototarazo genetika momban’ny mponina, dia hita fa azo itokisana ny fikarohana natao momban’ny firafitry ny mponina.
4. Hevi-dehibe — Ny fahasamihafana ara-genetika ao amin’ny *Loudetia simplex* dia manohana ny fisian’ny hivoka na kijana midadasika eto Madagasikara talohan’ny vanim-potoanan’ny fiovana paleoclimatika nisongadina. Na izany aza, ny fombam-pikarohana nampiasana teto dia tsy nahavita nanavaka ny fiovan’ny paleoclimatika akaikin’ny vanim-potoana lehibe nangatsiaka farany sy ny vokatry ny fitrandrahana nataon’ny olombelona. Mety ho fitaovana manan-danja amin’ny famakafakana ny firafitry ny mponina ny antotan-kevitra nampitombona na dia tsy misy fitaovana genomika iangaina aza.

**Fanambarana fiantraika ara-tsosialy:** Ny fanekena fa niely patrana ny hivoka itoeran’ny *Loudetia* talohan’ny fanjanahan’ny olombelona dia manamarika fa ireo hivoka ireo dia singa manan-danja amin’ny zavamananaina eto Madagasikara. Ilaina maika ny fahafantarana ara biolojika sy taxononomique ny zavamaniry sy ny biby amin’ny hivoka mba hanombanana ny loza mety hitranga amin’ny hivoka izay tsy ananana fahalalana maro sady heverina ho tany maina. Misy sakana ara-bola sy ara-pitaovana amin’ny fampiharana ny fandalinana momba ny fiarovana izay nampiasana fitaovana génomika ankehitriny. Nohatsarainay ny sasany amin’ireo fanamby mifandraika amin’ny famakafakana ara-genetika ny mponina manana ploidy maro tsy modely, izay tsy misy fitaovana genomika iaingana, amin’ny alàlan’ny fampivoarana loharanon-kevitra kajy mba hitrandrahana paikady izay mety hiteraka angon-drakitra mahomby tsy mitaky fahalalana mahakasika ny fototarazo ara-genetika ny zava-maniry izay tiana karohina.

## Introduction

The degree to which anthropogenic activities have transformed the ancestral vegetative composition of Madagascar’s ecosystems is heavily debated. While the narrative that most of central Madagascar was once covered in closed-canopy forests has been rejected (Bond et al., 2008; Vorontsova et al., 2016; Hackel et al., 2018; Solofondranohatra et al., 2020; Bond et al., 2022; Lehmann et al., 2022), the degree to which a former mosaic of grassland, shrubland, and forests has become more dominated by grasses is contentious (reviewed in Joseph et al., 2021). Understanding population dynamics of individual ecologically important species and plant communities will have profound effects on conservation policy decisions and the economic productivity of grasslands for local communities. However, it is difficult to determine where grasslands, which we consider to be an ecosystem where the ground layer is a majority of tussock-forming grass with little tree cover that experiences a dry season with regular fire regimes (e.g. Dixon et al., 2014), were in the past and how extensive they were. Stratigraphic sediment records from Central Madagascar indicate an increased relative abundance of grasses and fire through the Holocene (Burney, 1987; Gasse & Van Campo, 1998; Virah-Swamy et al., 2010), especially near the expected age of human colonization [c. two thousand years ago (ka); Dewar & Wright, 1993; Dewar et al., 2013; Pierron et al., 2017]. Megafaunal isotope records also support Holocene expansions of C_4_ grasslands but show a vegetative mosaic of C_4_ grasses and C_3_ vegetation was at least present (Crowley et al., 2021) near the last glacial maximum (LGM; c. 26.5-19 ka). Reconstructing spatio-temporal vegetative dynamics beyond the LGM begin to approach limitations of paleoecological data, but molecular evolutionary approaches can fill the gaps by providing probabilistic inferences through time.

Population genetic approaches for reconstruct demographic histories has been successfully applied to lemurs to reconstruct patterns of past forest connectivity (Quéméré et al., 2012; Yoder et al., 2016; Salmona et al., 2017; Teixeira et al., 2021; Tiley et al., 2022) and may similarly provide insights into the Late Pleistocene dynamics of grasses between the LGM and Last Interglacial period (LIG; c. 132-112 ka) or older. This strategy has been used to reconstruct vegetative dynamics of temperate Japanese grasslands from forb species (Yamaura et al., 2019).

An array of tools now exists to estimate the effective population size (*N_e_*) of species through time. As generating genomic data for non-model groups has become more accessible, population genomic methods that utilize precisely estimated site frequency spectra (SFS; Nielsen et al., 2000; Gutenkunst et al., 2009; Excoffier et al., 2013; Liu & Fu, 2020; Blischack et al., 2022) have become a valuable tool for reconstructing the demographic history of natural populations. There are still barriers to generating population genomic data though, either due to the lack of suitable tissue, an appropriate reference genome, or costs. Target-enrichment libraries have been transformative for the analysis of plant phylogenetic relationships (e.g. Breinholt et al., 2021; Baker et al., 2022), but the population genetic utility of such data is not well-explored. A survey of taxa with low sample sizes suggested population genetic parameters estimated from target-enrichment data could be unreliable, in addition to violating assumptions of the neutral coalescent model that underpins most tests of selection and demographic change (Slimp et al., 2021). However, the relatively long loci recovered by target-enrichment (or HybSeq) methods could be appropriate for a demographic modeling approach, the extended Bayesian skyline plot (EBSP; Heled & Drummond, 2008), which has been successfully applied to a range of conservation genetic questions despite some model violations. We are interested in exploring the potential of target-enrichment data for population or conservation genetic insights from plant groups that may be lacking genomic resources.

Here, we examined the population genetics of the grass species *Loudetia simplex* (Tristachyideae: Panicoideae: Poaceae). It is largely dominant in Madagascar’s Central Highlands (Koechlin, 1993; Hagl et al., 2021; Fig. 1a) and is notable for its fire- and grazing-adapted functional traits (Solofondranohatra et al., 2018). Population genetic analyses of microsatellites suggest Malagasy *L. simplex* has been isolated from mainland African populations with no detectable level of gene flow, and that there is additional population structure between the northern and southern extents of the species range across the Central Highlands of Madagascar (Hagl et al., 2021). Here, we use a subset of individuals analyzed by Hagl et al. (2021) along with a new sample from the center of the *L. simplex* distribution to explore the population genetic utility of target-enrichment data and its implications for the natural history of Madagascar’s grasslands. Because Malagasy *L. simplex* are putative polyploids (tetraploids and hexaploids; Hagl et al., 2021), we developed novel bioinformatic tools for the processing and analysis of polyploid data, integrated into the PATÉ allele phasing pipeline (Tiley et al., 2021).

**Figure 1.**
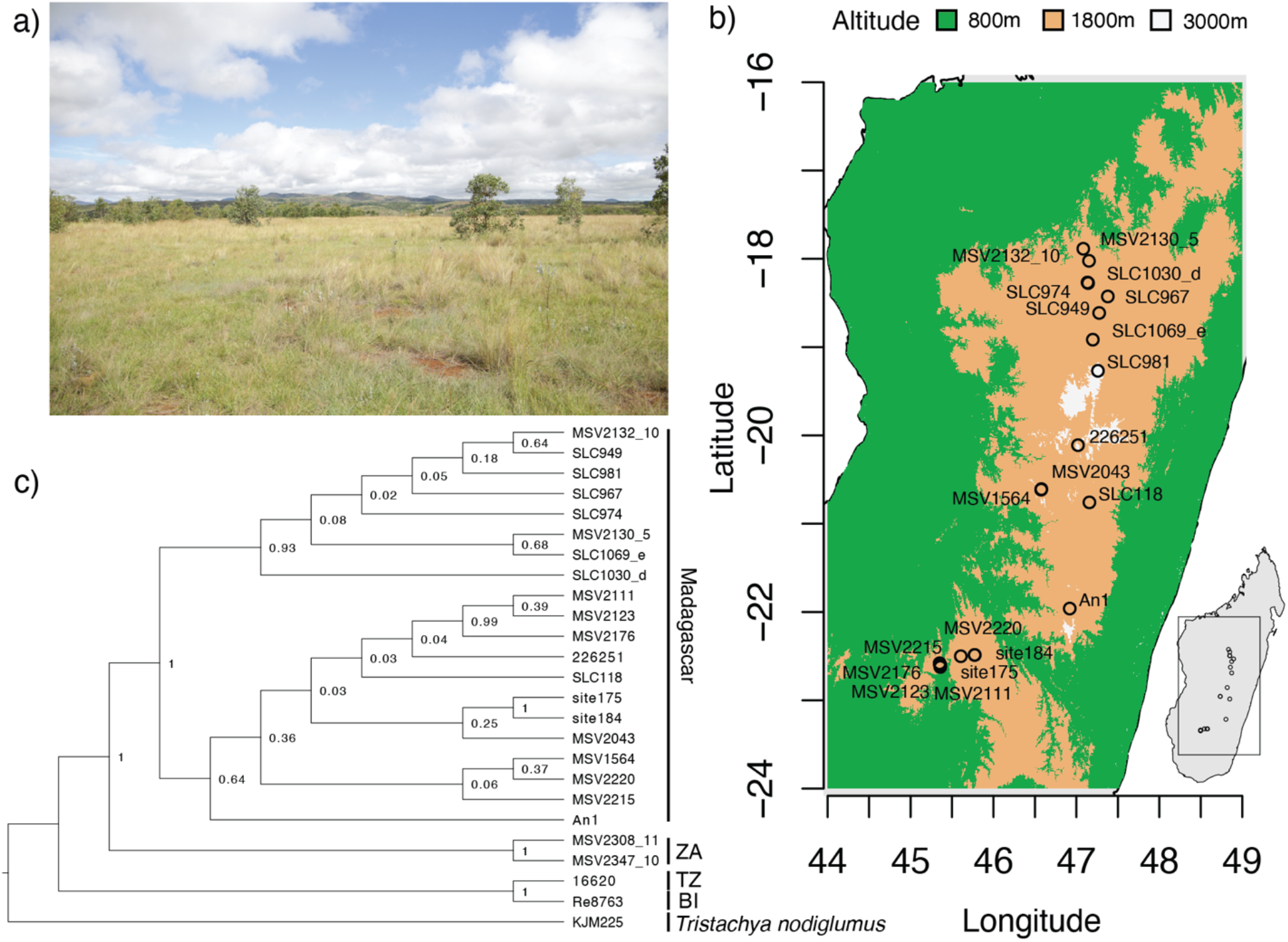
— *Loudetia simplex* distribution and single Madagascar origin. a) A field of *L. simplex* near Antsirabe Madagascar planted with eucalyptus (photo credit: George P. Tiley). b) Our sampling of *L. simplex* was based on collections made throughout Madagascar’s Central Highland Plateau, which is typically considered an altitude between 800 and 1800 meters. c) Phylogenetic relationships of individuals based on the phased (PH) data. Numbers next to nodes are posterior probabilities. ZA = South Africa, TZ = Tanzania, BI = Burundi.

## Materials and Methods

### Sampling and Sequencing

Target-enrichment data was generated for 24 individuals of *Loudetia simplex* and one outgroup, *Tristachya nodiglumus* (Supplementary Table S1). Twenty individuals were selected to best represent Madagascar from available collections as a single unstructured population and avoid complications of population substructure and low sample sizes in downstream analyses (Fig. 1b). Genomic DNA samples were used from Hagl et al. (2021) while new extracts used a modified CTAB protocol (Doyle & Doyle, 1987) from silica-dried leaf tissue. Genomic DNA was then sent to Rapid Genomics (Gainesville, FL, USA) where sample enrichment and sequencing was performed with a bait kit developed for angiosperms following Breinholt et al. (2021). All sample metadata is provided in Supplementary Table S1.

### Sequence Assembly and Processing

Initial contigs were assembled with HybPiper (Johnson et al., 2016) with default settings. We then used HybPiper’s supercontigs as reference sequences for PATÉ (Tiley et al., 2021; Crowl et al., 2022) to genotype individuals, filter low-quality variants with GATK v4.2.0 (McKenna et al., 2010; DePristo et al., 2011), and phase haplotype sequences using HPOP-G (Xie et al., 2016). Ploidy information for individuals (Supplementary Table S1) was based on previous microsatellite allele count data (Hagl et al., 2021); however, we also implemented a statistical test to infer ploidy level in PATÉ based on mixtures of distributions from allele balance data, similar to other approaches that use the number of reads supporting an alternate allele over total reads to infer ploidy (Weiß et al., 2018). We extended the mixture models of Tiley et al. (2018) to determine the ploidy of a sample with the model weights as a selection criterion (Supplementary Methods: *Ploidy Test*). PATÉ control files with all filtering options for GATK are provided in the Supplementary Data (https://doi.org/10.5061/dryad.905qfttqx). Because target-enrichment probes may represent multiple exons from the same gene with overlapping flanking regions, we were concerned that some assembled loci might overlap and violate some independence assumptions in downstream analyses. Therefore, we performed an all-by-all blastn search (Camacho et al., 2009) with an e-value of 1 × 10^−5^ among the supercontig assemblies followed by single-linkage clustering (https://github.com/gtiley/HybSeq-Cleaning).

Only the longest locus per cluster was retained for downstream analyses.

To investigate nucleotide variation and the potential population genetic value of non-coding versus coding regions of target-enrichment loci, we identified the following regions:

1. *Left Flank* – Non-coding sequence upstream of the 5’ start codon or splice site.
2. *Core* – The coding region targeted by probes for sample enrichment.
3. *Right Flank* – Non-coding sequence downstream of the 3’ stop codon or splice site.

Identification of individual regions was automated (https://github.com/gtiley/Locus-Splitter) based on the tblasx program from BLAST v2.10.0 (Camacho et al., 2009) to find in-frame alignments between our assembled sequences and the original transcriptome loci used for probe development. Individual regions were aligned with MAFFT v7.471 (Katoh & Standley, 2013). The aligned regions were then concatenated together to make whole-locus alignments.

Because we are interested in investigating the analytic impact of ploidy in sequence data, we created three datasets from the phased ploidy-aware data:

1. *One Haplotype (OH)* – Only one phased haplotype sequence was randomly selected from each individual for each locus.
2. *Ambiguous Genotype (AG)* – The genotypic information for an individual is represented as a single sequence with IUPAC ambiguity codes for biallelic sites. Analyses that treat IUPAC codes as missing data were randomly resolved into two sequences.
3. *Phased Haplotypes (PH)* – All phased haplotype sequences for an individual are used.

We expect the OH data to be representative of majority consensus sequences (e.g. HybPiper supercontigs) since our target-enrichment loci are anchored in single exons and we anticipate that any risk of assembling chimeric sequences from such data without paralog warnings is low. Any site genotyped as polyallelic, having more than one alternate allele, was assumed to be an error and treated as missing sequence across all data sets. All expected copies of a locus are represented even if they are invariant. For example, some individuals in the PH data may have six identical sequences at a locus if they are hexaploids and every site is homozygous. While the ploidy at an individual locus may vary due to differential gene loss among chromosomes, this is a simplifying assumption to preserve expected allele frequency information.

### Phylogeny Estimation

A species tree was estimated for all *L. simplex* individuals using whole-locus sequences for the OH, AG, and PH datasets with BPP v4.6.2 (Flouri et al., 2020). We chose a full-likelihood multispecies coalescent (MSC; Rannala & Yang, 2003) model since it is robust to low-information gene tree estimates (Xu & Yang, 2016). Markov chain Monte Carlo (MCMC) analyses collected 10,000 posterior samples, saving every 10 samples after a 10,000 generation burnin. Model and prior specifications are provided in the Supplementary Material (Supplementary Methods: *BPP Prior Specification*). The starting tree for our Bayesian species tree analyses was based on a concatenated maximum likelihood (ML) estimate from the OH data with IQTREE v2.1.2 (Minh et al., 2020). We used unpartitioned analyses with ModelFinderPlus (Kalyaanamoorthy et al., 2017) for the OH, AG, and PH data to estimate ML evolutionary distances between individuals, with random concatenation of haplotypes within individuals.

### Population Structure

Potential population structure within our sample of *L. simplex* from Madagascar was explored with STRUCTURE v2.3.4 (Pritchard et al., 2000). For each dataset, we sampled only one biallelic SNP per locus, if present, and required a minimum minor allele count of three with no more than 25% missing data. To explore uncertainty due to sampling error of sites in a target-enrichment dataset that is relatively small compared to contemporary population genomic data, we generated 10 datasets with random SNP sampling. The number of clusters that best satisfy Hardy-Weinberg assumptions was determined with the 𝛥𝐾 method (Evanno et al., 2005).

Principal component analyses (PCAs) were conducted on the OH, AG, and PH datasets with adegenet v2.1.3 (Jombart & Ahmed, 2011) by compressing allele counts to the same dimension as the lowest ploidy level present. For example, a hexaploid with three reference and three alternate alleles would be scored as a 2 on a scale from 0 to 4, to be comparable to tetraploids in the dataset. Additional details on allelic imbalances would be lost as, for example, hexaploids with either one or two alternate alleles would be represented as a 1 on the tetraploid scale. Scripts for generating PCA or other clustering-ready matrices from fasta files are available on GPT’s GitHub (https://github.com/gtiley/fasta-conversion). Additional details about compression of PCA data matrices can be found in the Supplementary Material (Supplementary Methods: *PCA Input Data*). The presence of isolation-by-distance as an explanation for observed patterns of genetic structure and ordination was tested with the patristic distances from the ML analyses or the uncorrected pairwise distances (p-distances) regressed against the natural logarithm of the great circle distance between individuals. A Mantel test was performed for isolation-by-distance analyses in the ape v5.6.2 R package (Paradis & Schliep, 2019).

### Population Genetic Summary Statistics

The pairwise nucleotide diversity (𝜋), number of segregating sites (*S*), and Tajima’s *D* were calculated across locus regions and datasets with the PopGenome v2.7.5 R package (Pfeifer et al., 2014). The significance of Tajima’s *D* statistics was determined by 100 standard coalescent simulations with ms (Hudson, 2002). We generated 200 bootstrap replicates of each alignment to calculate a percentile bootstrap 95% confidence interval and compare the relative estimation error between regions and datasets using the root mean square error (RMSE) from the bootstrap samples. The scaling and interpretation of parameters are described in the Supplementary Material (Supplementary Methods: *Summary Statistics*). We tested for differences in the distribution of summary statistics among locus regions (left flanks, cores, and right flanks) and data types (OH, AG, and PH) with Wilcoxon Rank-Sum tests. We tested for biases in Tajima’s *D* results among locus regions and data types with 𝛸^#^ tests.

### Demographic History

We chose the extended Bayesian skyline plot (EBSP, Heled & Drummond, 2008) as the most appropriate model of *N_e_* change through time for our sample of target enrichment data. Here we assumed a strict molecular clock underlying all lineages but allowed rates of evolution to vary among loci. One locus was fixed to the clock rate and all other locus rates were relative. Rates of molecular evolution among plants are especially complicated, as there can be life histories with variable reproductive strategies, unknown generation times, and overlapping generations. Thus, we relied on an estimate of the neutral substitution rate at third-position synonymous sites from grass genomes (Christin et al., 2014) and constructed a vague prior around this mean rate, such that any biologically plausible uncertainty in the rate of evolution is reflected in our posterior sample. It is a tenuous rate estimate at best, since it remains unknown whether polyploid *L. simplex* largely propagates clonally, through apomixis, or through sexual reproduction. The MCMC analyses for the OH, AG, and PH data collected 20,000 samples after a burnin of 50,000,000 and sample interval of 10,000. Two replicates were run for each analysis to evaluate convergence and mixing was assessed with Tracer v1.7 (Rambaut et al., 2017). All model and prior specifications are provided in the Supplementary Material (Supplementary Methods: *EBSP Prior Specification*) and are replicable from XML files on Dryad (https://doi.org/10.5061/dryad.905qfttqx).

### Adequacy of Demographic Models

To explore if EBSP models could reliably differentiate between Late Pleistocene versus Holocene population size increases, we evaluated the accuracy of EBSP inferences with simulations that captured some features of our target-enrichment data. We simulated 57 loci that were 1000 bp in length for 20 haploid individuals 100 times with ms (Hudson, 2002) and seq-gen (Rambaut & Grass, 1997) for four scenarios: 1) a constant *N_e_* of 100,000, 2) an *N_e_* increase from 10,000 to 100,000 individuals roughly coinciding with the LGM at 20,000 years in the past, 3) the same ten-fold increase but at 2,000 years in the past to reflect anthropogenic expansion, and 4) a scenario combining both LGM and anthropogenic effects by exponentially increasing *N_e_* from 10,000 to 100,000 between 20,000 years ago and the present. Each simulated dataset was then evaluated with the same EBSP model using a fixed clock used for our empirical analyses, except the MCMC algorithm collected 10,000 posterior samples with a sample interval of 1,000 after a 100,000 sample burnin. Commands for simulations are given in the Supplementary Material (Supplementary Methods: *Simulating Demographic Histories*).

## Results

### Sequencing Statistics

The probe set targeted 408 loci from which an average of 327 loci were assembled by HybPiper (Supplementary Table S2). These were supported by an average of 217 read pairs per locus. Although there was variable read depth across loci and individuals, the number of assembled loci was significantly correlated with read depth (Pearson’s correlation coefficient = 0.57, *p*-value = 0.002). After joint-genotyping all individuals as diploids, normal mixture models suggested all individuals were hexaploids, with the exception of two individuals from South Africa (Supplementary Table S3). Some individuals were inferred as pentaploids, but since this was unexpected based on previous observations from microsatellites (Supplementary Table S1) these cases were assumed to be hexaploids. Although available evidence leaves the ploidy of some individuals ambiguous and raises skepticism of the inference methods, genotyping and phasing of individuals proceeded with the ploidy suggested by our mixture models. We inferred an average of 247 loci with allelic variation, which had a heterozygosity of 0.012 variants per base. After reducing the number of loci with BLAST clustering to avoid tight linkage, a final 301 loci were retained from the OH, AG, and PH datasets for downstream analyses. The average number of haplotype blocks across all loci was 1.005, meaning that in very few cases there was insufficient data to phase across the entire locus (Supplementary Table S2). In cases where multiple variants within a locus could not be phased, only the variants from the longest block were retained and variants on the shorter block were replaced with missing data.

### Phylogeny Estimation

Species trees revealed all Malagasy *L. simplex* form a well-supported clade (Supplementary Fig. S1), but there was disagreement in the biogeographic history within Madagascar based on the treatment of ploidy. Individuals in northern Madagascar formed a clade with a posterior probability of 0.9 or greater across the OH, AG, and PH analyses. However, the individuals from central and southern Madagascar were paraphyletic in the OH and AG analyses (Supplementary Figs. S1a and S1b). A clade of central and southern individuals was recovered in the PH analysis, but with a weak posterior probability of 0.64 (Fig. 1c; Supplementary Fig. S1c). Individuals from the center of the *L. simplex* distribution appear to be the earliest diverging lineages in the OH and AG analyses (Supplementary Figs. S1a and S1b), or with respect to the sample of central and southern individuals in the PH analysis (Fig. 1c; Supplementary Fig. S1c).

### Population Sub-Structure in Madagascar

STRUCTURE analyses recovered a variable number of Hardy-Weinberg groups across the ten datasets that randomly sampled one non-singleton biallelic SNP per locus (Supplementary Table S4). For the OH data, we found a *K* of two, three, or four in eight, one, and one of the respective replicates. For AG data, the optimal number of clusters was more ambiguous with a *K* of two, three, and four recovered for four, five, and one replicates. PH data found *K* of two and three for eight and two replicates. We note that the method of Evanno et al. (2005) cannot infer if one panmictic population is the best model, and while there is reasonable geographic structure at *K* of two weighted across biological and technical replicates (Fig. 2a) the non-zero admixture between individuals at the most extreme ends could indicate a single population with a cline of genetic variation. This is further supported by the lack of distinct cluster assignments at higher values of *K* despite some evidence for additional population structure (Supplementary Fig. S2). A weak signal of isolation-by-distance was detected when regressing genetic distance against spatial distance (Fig. 2b). Notably, the AG data inferred much higher pairwise distances between individuals; although, the relative differences and overall trend were similar. The same patterns can be observed when using patristic distances from ML phylogenetic estimates, although the disparity between AG versus OH or PH distances is reduced, as is the percent of genetic variation explained by spatial distances (Supplementary Fig. S3).

**Figure 2.**
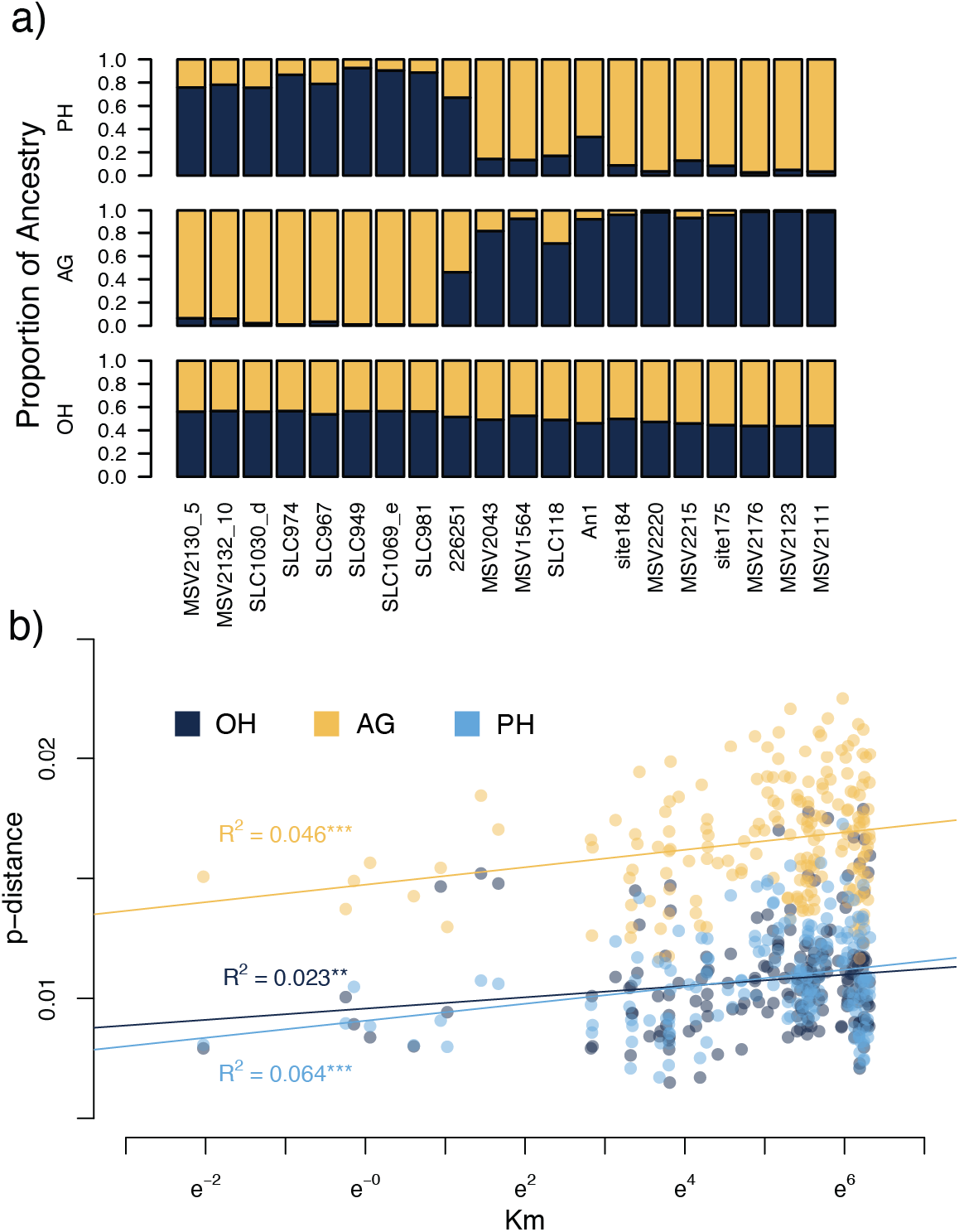
— Patterns of genetic structure and variation in Madagascar. a) Aggregated STRUCTURE results across runs for an *a priori* number of clusters of two (*K* = 2) for the different data types. c) A plot of natural-log-scaled geographic distance against the pairwise distance (p-distance). Lines are from a least-squares linear regression with the coefficient of determination (R^2^) shown for each data type. Statistical significance was determined with a Mantel test using 9999 permutations. Significance levels are *p*-value < 0.05 (*), *p*-value < 0.01 (**), and *p*-value < 0.001 (***). Colors represent different data types: one haplotype (OH), ambiguous genotypes (AG), and phased haplotypes (PH).

PCAs reflected findings from STRUCTURE analyses and reinforce the presence of a genetic cline (Fig. 3). For OH data, where only one reference or alternate allele was observed per individual, Malagasy *L. simplex* is clearly distinguished from mainland African individuals (Fig. 3a), but there is no clear separation of individuals based on geographic location (Fig. 3b). PCAs of data that accounts for heterozygosity but not dosage recovered a single Malagasy versus mainland African pattern (Fig. 3c), PCAs reflect a north-to-south gradient of genetic variation regardless of whether allelic dosage was treated as hexaploid (Fig. 3d), tetraploid (Fig. 3e), or diploid (Fig. 3f). The total percent of variance explained by principal component axis 1 (PC1) and axis 2 (PC2) increased as allelic dosage was compressed from a hexaploid to a diploid state.

**Figure 3.**
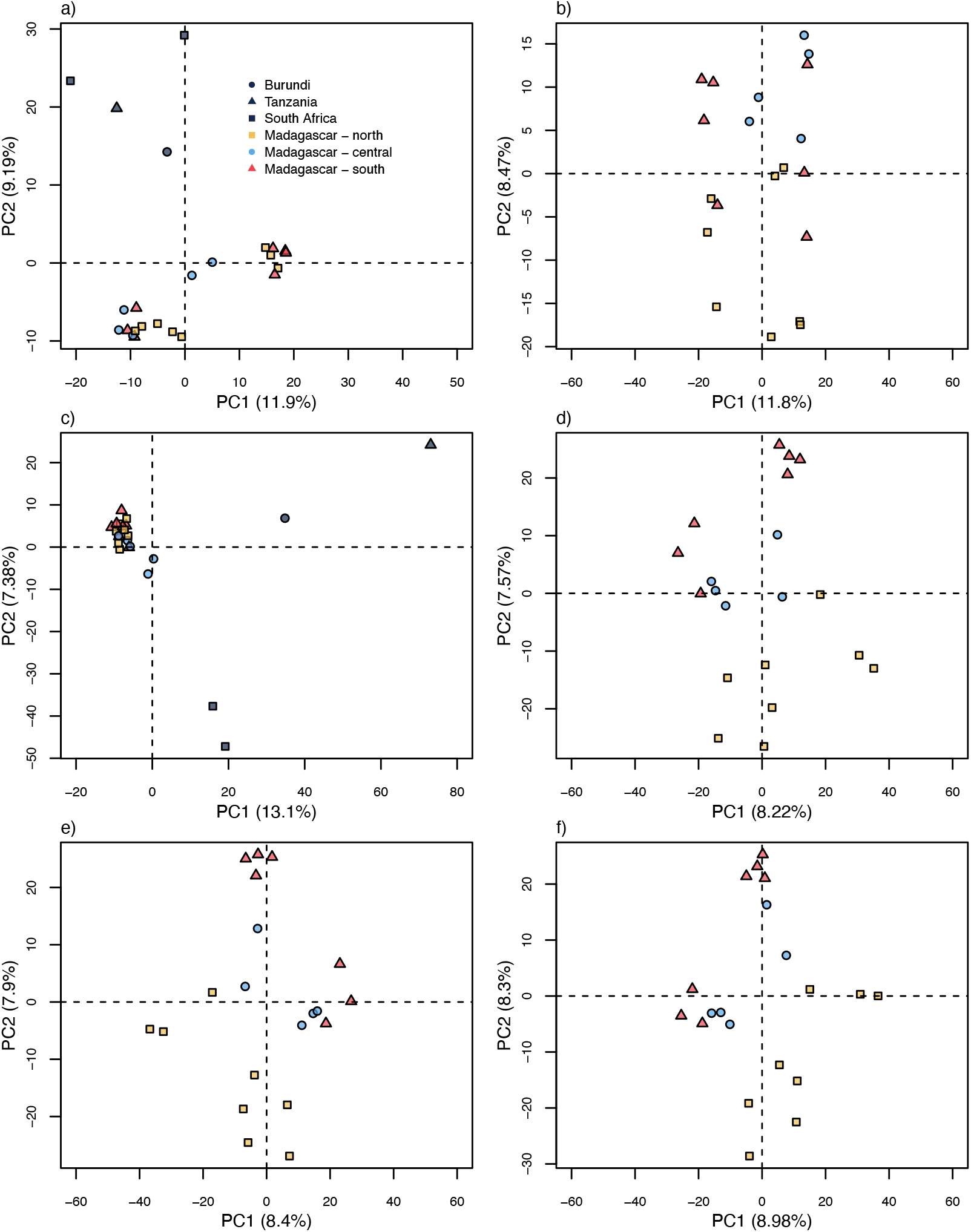
— Principal component analyses across data types. a) All individuals using haploid (OH) data. b) Only Malagasy individuals using OH data. c) All individuals treating all sites as diploid. d) Only Malagasy individuals treating all sites as hexaploid. e) Only Malagasy individuals compressing unbalanced allelic dosage to single categories. f) Only Malagasy individuals treating all sites as diploid.

### Variation in Summary Statistics

There were significant differences in the distributions of 𝜋, *S*, and Tajima’s *D* among locus regions and between the OH, AG, and PH data (Fig. 4). For 𝜋, there was always a difference between flanking regions and the exon core (Supplementary Table S5), with the cores surprisingly having more variation. Differences were also observed between the flanking regions and cores for *S* and Tajima’s *D*, except when comparing the left flanks and cores for OH data (Supplementary Table S5). The only case of a difference between the left and right flanks was when estimating 𝜋 for AG data (Supplementary Table S5). The OH and PH data resulted in similar estimates of 𝜋; however, the AG and PH data had similar distributions of *S* (Supplementary Table S5). Any differences in Tajima’s *D* statistics were largely explained by an over-representation of values less than negative two, indicating positive selection or population size expansion, in the OH data and among core regions across datasets (Table 1).

**Figure 4.**
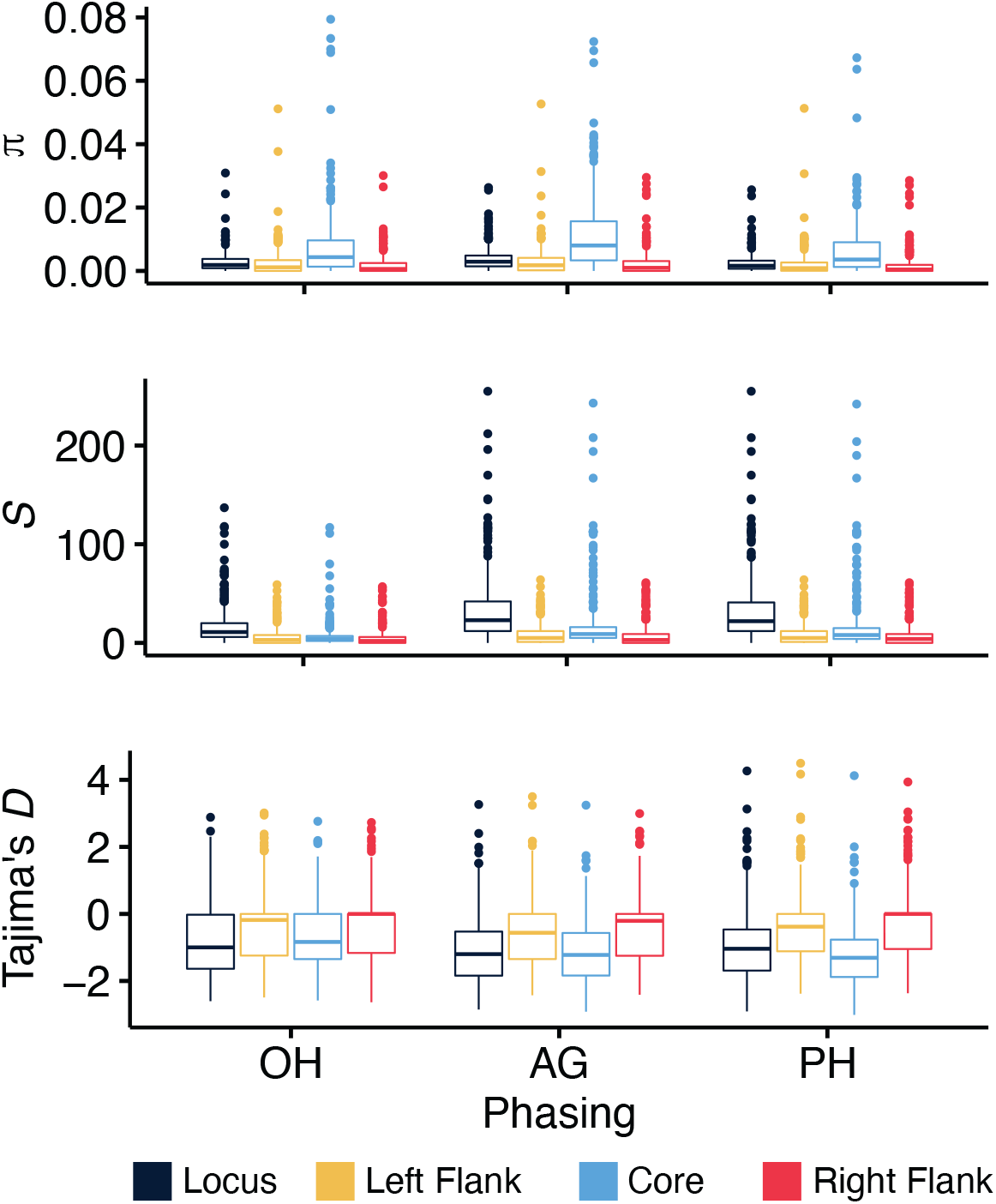
— Distributions of summary statistics across locus regions. Box-and-whisker plots show distribution medians and quartiles for the expected pairwise nucleotide divergence (𝜋), the number of segregating sites (*S*), and a normalized measure of the difference between two estimators of nucleotide divergence, indicating deviations from neutral expectations (Tajima’s *D*).

**Table 1.** 𝛸^2^ test result from Tajima’s *D* statistics across datasets and locus regions.

| Tested Effect | Fixed Effect | $\chi^2$ | $p$-value | Largest Residual |
| --- | --- | --- | --- | --- |
| Region | OH | 7.72 | 0.27 | Under-representation of Tajima's $D < -2$ in core regions |
| | AG | 37.77 | 0.0001 | Over-representation of Tajima's $D < -2$ in core regions |
| | PH | 83.47 | 0.0001 | Over-representation of Tajima's $D < -2$ in core regions |
| Dataset | Left Flank | 12.65 | 0.046 | Over-representation of Tajima's $D < -2$ in OH dataset |
| | Right Flank | 8.61 | 0.19 | Over-representation of Tajima's $D < -2$ in OH dataset |
| | Core | 57.15 | 0.0001 | Over-representation of Tajima's $D < -2$ in OH dataset |

We attempted to explore biases in summary statistic estimates due to sampling error of sites through bootstrapping the OH (Supplementary Fig. S4), AG (Supplementary Fig. S5), and PH data (Supplementary Fig. S6). Our observed summary statistics typically fell outside of their percentile bootstrap 95% confidence intervals, but other confidence interval approximation techniques (e.g. Efron & Narasimhan, 2020) may be more appropriate for population genetic summary statistics (Leblois et al., 2003). However, we used the bias, difference between the bootstrap mean and observed estimate, and the RMSE of bootstrap replicates with respect to the observed estimate, to gain some general insights. Notably, the bias of 𝜋 is larger and much noisier in the AG data (Supplementary Fig. S5) compared to the OH (Supplementary Fig. S4) and PH (Supplementary Fig. S6) data. Even if the bias for a whole-locus is low, there can be high variability between regions. For the observed values, estimates are not always proportional among regions either. The RMSE of an estimate tends to decrease along with genetic variation, implying some role of genotyping error or rare variants and singletons inflating values at the higher end of the spectrum. Loci with very low levels of genetic variation can also be problematic though, as the RMSE for flanking regions increases towards the tail of the distribution, creating opportunity for sampling error to wildly swing estimates regardless of ploidy (Supplementary Figs. S4-S6). The number of segregating sites increased with ploidy of the data, but 𝜋 was relatively stable. However, differences between 𝜋 and *S* will affect estimates of Tajima’s *D*, and this could affect interpretations of demography or selection acting on loci; although, the majority of loci and their regions were not significantly different from neutral expectations given the simulation-based test statistics (Supplementary Table S6).

### Population Size Change Through Time

After filtering for longer loci with low amounts of missing data, 57 loci were retained for EBSP analyses. Reducing our data to a subset of more informative loci was done for computational convenience, because EBSP models are parameter-rich and require long MCMC runs for convergence. Runs converged for each data type for both analyses with a fixed rate of molecular evolution (Fig. 5) and variable rate (Supplementary Fig. S7). All analyses suggested the *L. simplex N_e_* increased through the Late Pleistocene, well before the LIG, approximately one million years ago. Between the LGM and period of potential human colonization of Madagascar in the Holocene, *N_e_* had already reached present-day estimates. *N_e_* estimates through the Holocene were only possible for the PH data, as accounting for the within-individual variation should have captured more recent coalescences. Simulations cast doubt on the seriousness with which EBSP results can be interpreted through the Holocene though. For example, when *N_e_* was truly constant through time, some individual EBSP analyses would suggest recent *N_e_* increases; although, the mean estimate across simulations recovered the constant population scenario (Fig. 6a). This problem was exaggerated in the presence of model over-fitting (Supplementary Fig. S8). In scenarios with recent *N_e_* increases, the contemporary *N_e_* was underestimated but a trend indicating a recent *N_e_* increase was detectable (Figs. 6b-6d). However, simulations indicated it would not be possible to differentiate between *N_e_* increases that occurred during the LGM (Fig. 6b) versus the Late Holocene (Fig. 6c), let alone some combination of paleoclimate and anthropogenic effects (Fig. 6d). Interpreting these recent *N_e_* increases from EBSP analyses becomes impossible with model over-fitting (Supplementary Fig. S8).

**Figure 5.**
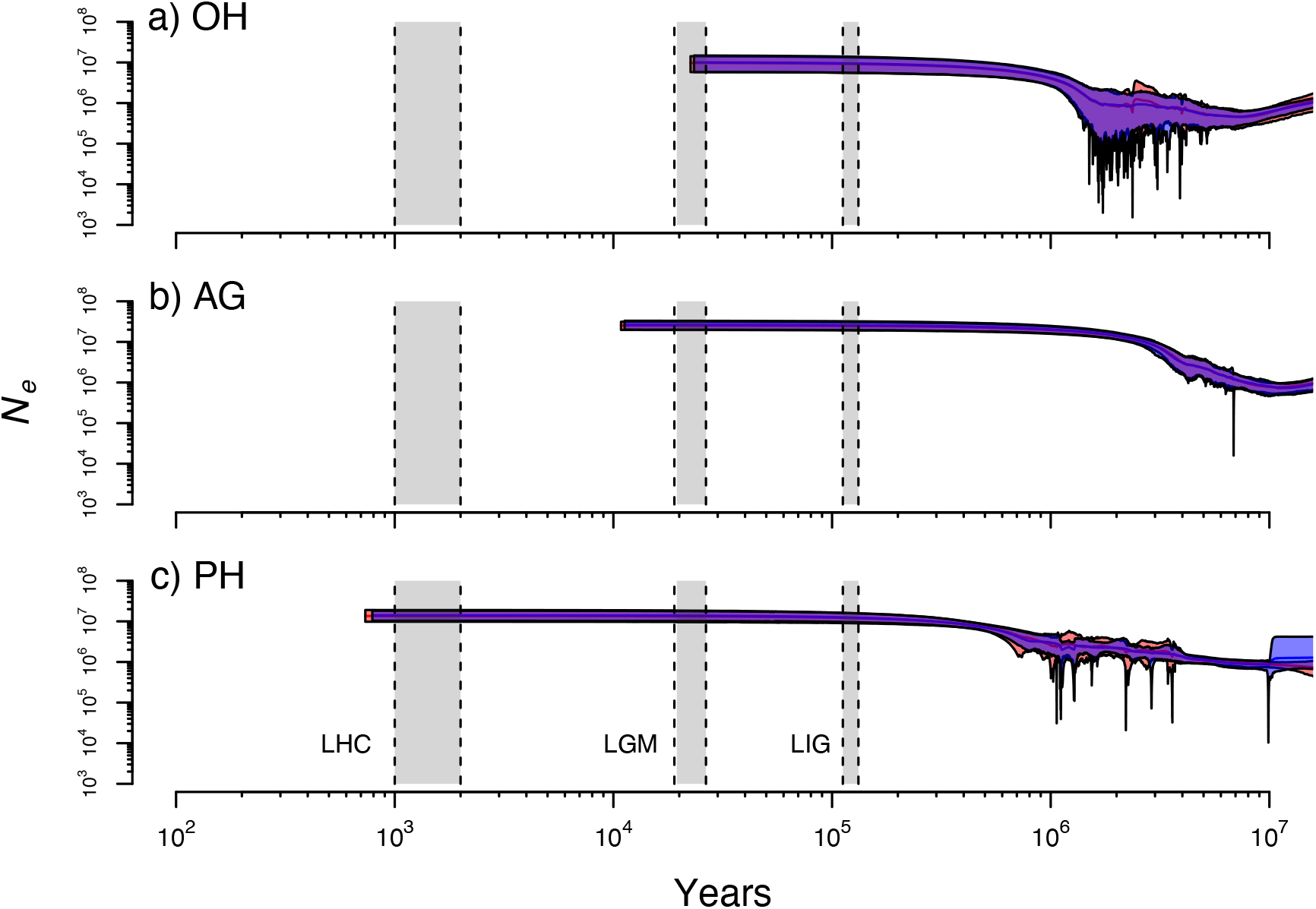
— Extended Bayesian Skyline Plot results with fixed clock rates. Two independent runs are shown with mean population sizes shown by solid lines. Polygons show the 95% HPD intervals with the overlap between runs in purple. Results are shown for the a) OH, b) AG, and c) PH data. Relevant time periods are shown in gray, which includes likely human colonization (LHC), the last glacial maximum (LGM), and last interglacial period (LIG).

**Figure 6.**
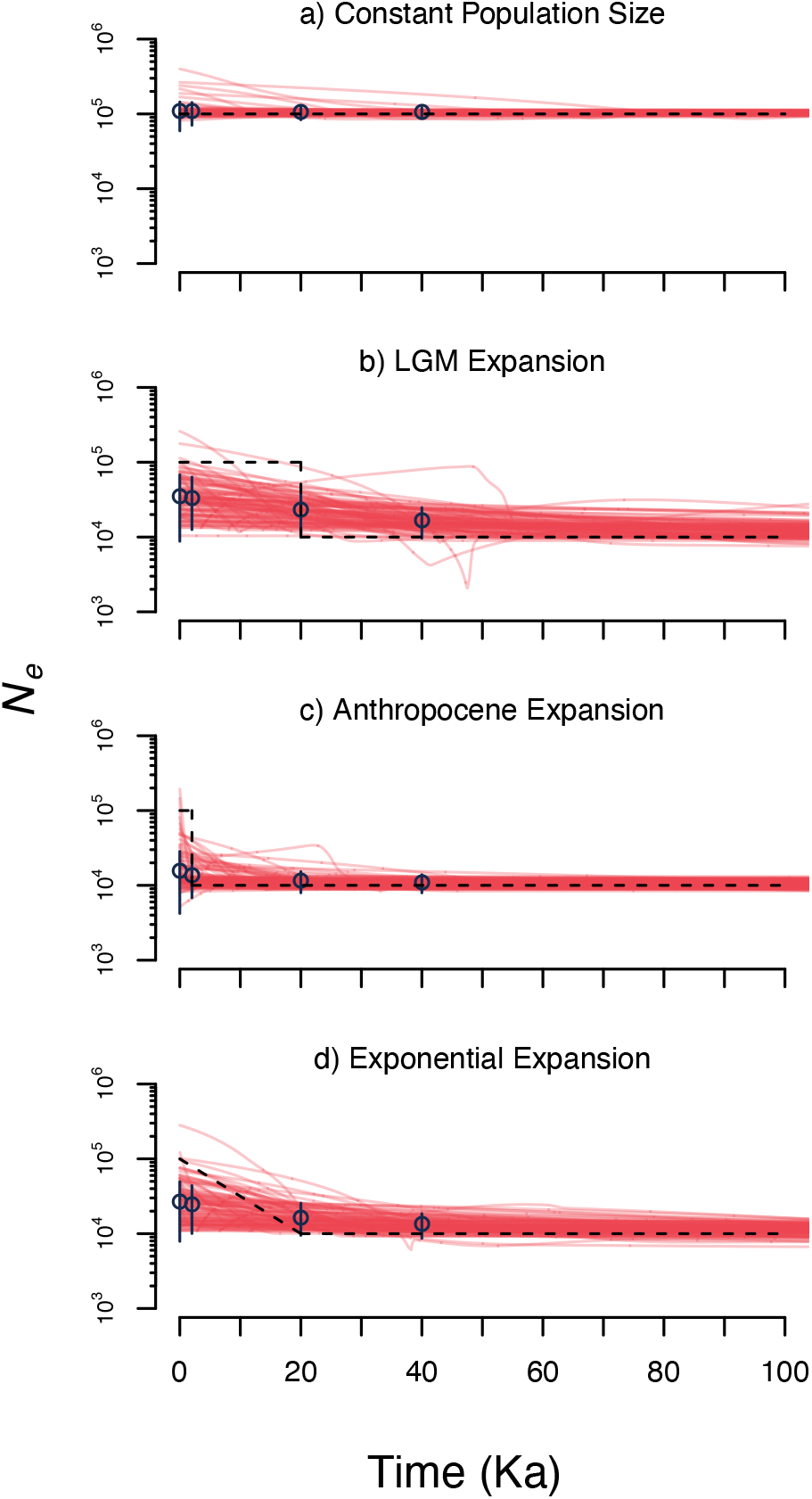
— Simulation results from EBSP analyses for an appropriate number of change points. Individual red lines are the mean *N_e_* estimates from 100 simulated data sets. Blue points and vertical lines are the mean of mean *N_e_* estimates and their 95% HPD intervals. The dashed black line is the true demographic history of the population.

## Discussion

### Robustness of Analyses Across Loci and Ploidy

Although target-enrichment data has been transformative for species-level phylogenomic studies (Breinholt et al., 2021; Baker et al., 2022), the robustness of population genetic estimates for a sample of closely related individuals was largely unexplored (but see Slimp et al., 2021). Some of the most important measures of genetic variation for populations were susceptible to sampling error, especially in the flanking intron regions (Supplementary Figs. S4-S6). While estimates from cores showed less error, they will lead to some violations of population genetic methods that assume evolution consistent with neutral theory, given potential signatures of purifying selection (Table 1). This creates some conflict, since most of the information is contained within the exon cores of target-enrichment loci (Fig. 4), presumably due to read coverage. Because Tajima’s *D* alone cannot disentangle demography from selection, some judgment calls and tolerance for downstream model violations are necessary. The majority of loci examined were consistent with neutral expectations based on simulations (Supplementary Table S6), and the proportion of loci with significant Tajima’s *D* statistics were exaggerated when ignoring heterozygosity (Table 1). Although, the estimates of nucleotide diversity and simulations under the standard neutral coalescent used here should be reasonable for polyploids in cases of perfect multisomic inheritance in outcrossers (Arnold et al., 2012; Blischak et al., 2023), we note that the presence of subgenome structure and selfing could cause deviations in expectations of allele frequencies (e.g. Ronfort et al., 1998; Meirmans & Van Tienderen, 2013; Blischak et al., 2023). The mechanism of polyploidization is unknown in *L. simplex*, but our results do provide some expectations of how genotyping choices for polyploids can impact standard summary statistic estimates.

Predicting the effects of assumptions about genotype on estimates of genetic variation and downstream analyses are not always straightforward. For example, the pairwise distances (Fig. 2b) or evolutionary distances (Supplementary Fig. S3) between individuals was higher when assuming all individuals were diploid (AG) compared to assuming all individuals were haploid (OH) or genotyped at their anticipated ploidy level (PH). If we were willing to ignore the absolute estimates of genetic distance, the interpretation of isolation-by-distance analyses was the same across the OH, AG, and PH data. This implies that biologically meaningful conclusions can arise even with uncertainty in ploidy and phasing. Interestingly, treating all individuals as diploid was sufficient for STRUCTURE to recover northern and southern clusters with some admixed central individuals of varying proportions (Fig. 2a; Supplementary Fig. S2). Similarly, PCAs reflected the geographic distribution of our *L. simplex* samples in the AG and PH data with, surprisingly, the AG data explaining more of the observed genetic variance (Fig. 3f) than the PH data (Fig. 3d).

This may be due to genotyping error at higher ploidy levels caused by read stochasticity and a flat genotype likelihood surface (Blischak et al., 2016) or caused by incorrect ploidy inferences. A similar mixture model approach to the one implemented here (Weiß et al., 2018) has been shown to have good performance based on comparisons of allele balance distributions from target-enrichment data and flow cytometry-based ploidy estimates (Viruel et al., 2019). We are cautiously optimistic about the prospects of direct ploidy estimation from sequence data, because likelihood-based criteria can be prone to overfitting (Tiley et al., 2018). However, our ploidy estimates are generally consistent with the previous inferences from microsatellite allele counts (Hagl et al., 2021; Supplementary Table S1; Supplementary Table S3). A positive insight from our comparisons across ploidy levels is that if unknown individuals are genotyped as diploid, there will be sufficient information about population structure, and that ploidy estimation should not be a barrier to investigations of natural plant populations with little *a priori* cytological or genetic knowledge.

### Limits to Differentiating Anthropogenic and Paleoclimate N_e_ Change

Large population genomic studies in plants that can leverage demographic estimates from SFS (e.g. Excoffier et al., 2013; Liu & Fu, 2020; Blischak et al., 2023) are commonly focused on temperate regions (e.g. Yamaura et al., 2019) and species exploited for energy or agriculture (e.g. Huang et al., 2014; Wang et al., 2017). Thus, it is tempting to leverage models that use gene trees as data such as the EBSP (Heled & Drummond, 2008) or other extensions of skyline plots (Pybus et al., 2000), since they could be paired with relatively low-cost target-enrichment strategies that yield multi-locus nuclear data while stretching material quality.

While our analyses provide valuable insights into the ancient origins of *L. simplex* dominated grasslands, both our empirical and simulated analyses show we cannot differentiate between *N_e_* increases that occurred during human colonization or paleoclimate change near the LGM. The change in vegetative composition towards more C_4_ grass species in Central Madagascar is well-documented (Burney, 1987; Gasse & Van Campo, 1998; Virah-Swamy et al., 2010; Crowley et al., 2021). It is likely that these increases in C_4_ grasses included *L. simplex*; although, an increase in census size may not be reflected by *N_e_* (Fig. 5; Supplementary Fig. S7), the expected population size under a neutral coalescent model. Our simulations showed that detecting these recent increases is difficult, and that we cannot differentiate between a ten-fold increase that occurred 200 (2 ka) or 2000 generations ago (20 ka; Fig. 6). Diagnosing a true *N_e_* change becomes impossible in the presence of model overfitting (Supplementary Fig. S8) and diagnosing these problems in empirical data can be difficult aside from convergence assessments.

These limitations do not necessarily disagree with an assessment of EBSP models from a larger genomic dataset of endemic palm species in Madagascar (Helmstetter et al., 2021). Helmstetter et al. (2021) found that the estimated *N_e_* trends were valuable predictors of IUCN risk assessments, and that population decreases could be detected at anthropogenic time scales. However, it should be easier to detect population declines than population increases in a system that already had a large degree of standing genetic variation, such as *L. simplex*. This is because the time until the most recent common ancestor of a genealogy will be much more recent in a small population than a large one (e.g. see equation 2.13 from Gillespie, 2004). Analyses of demographic models with genomic SFS should be able to increase resolution at recent anthropogenic time scales (Patton et al., 2019) and differentiate anthropogenic from paleoclimatic demographic effects (Tiley et al., 2022), but this would require more extensive sampling and sequencing. Additionally, our analyses treat all individuals as a single population, and since we detected some level of population structure, we cannot rule out their confounding effects on analyses of *N_e_* through time (Heller et al., 2013; Mazet et al., 2016) or other confounding affects such as clonal growth and overlapping generations. Population-level sampling of multiple sites throughout a species range should be used to control for population structure, while accounting for localized effects such as agriculture and grazing versus regional climate effects.

### New Insights into Loudetia simplex Evolution and Policy Implications

Our analyses yielded new findings about the evolution of *L. simplex* that are potentially consequential for the broader understanding of Malagasy grasslands. First, our species trees (Supplementary Fig. S1) indicate that individuals from what we call Central Madagascar – a loose term meant only to indicate individuals from the center of the sampled *L. simplex* distribution – represent the ancestral distribution. It is likely that *L. simplex* dominated grasslands have origins in some of the higher elevation regions of the Central Highlands. For example, MSV1564 comes from the Itremo protected area (Fig. 1b; Supplementary Table S1), and further investigations of this region should be productive for characterizing natural grasslands in Madagascar. The hypothesis of Central Highland origins, which we consider 800m in altitude and above, is also supported by our analyses of population structure (Fig. 2a; Supplementary Fig. S2) and the weak but significant pattern of isolation-by-distance (Fig. 2b; Supplementary Fig. S3).

Our models of *N_e_* change through time suggests that *L. simplex* grasslands were already extensive (Fig. 5; Supplementary Fig. S7) well before any hypothesized human presence in Madagascar (c. 10 ka; Hansford et al., 2018) let alone human colonization (c. 2 ka; Dewar and Wright, 1993; Dewar et al., 2013; Pierron et al., 2017). We found an approximate ten-fold increase between one MA and the LIG which is beyond the limitations of available paleoecological records (Burney, 1987; Gasse & Van Campo, 1998; Virah-Swamy et al., 2010; Railsback et al., 2020; Crowley et al., 2021). Strong assumptions about the mutation rate and generation time to calibrate analyses to absolute time and *N_e_* were necessary, but our interpretations of results are within the biologically plausible uncertainty of these parameters. Even if the per-year substitution rate was overestimated by a factor of ten, the *N_e_* increase would still be constrained to the Late Pleistocene, occurring between the LIG and LGM. These analyses, especially in the context of a previous investigation of population structure (Hagl et al., 2021), suggest that *L. simplex* dominated grasslands have pre-human origins and likely experienced some *N_e_* increase coincident with Pleistocene climate change. Tree planting programs have been deployed to combat the devastating habitat loss and deforestation in Madagascar since the 1950s (Green & Sussman, 1990; Harper et al., 2007; Vieilledent et al., 2018; Schüßler et al., 2018), with country-wide forest loss from 1953 to 2014 at 44% (Vieilledent et al., 2018). The results of our study strongly imply that tree planting efforts should be cautious of treating *L. simplex* dominated grasslands as degraded forests, and we suggest that focusing resources on forest edges with native species will be more productive (Miandrimanana et al., 2019).

### Potential Use Cases of Target-Enrichment Data for Conservation Genetics

One application of target-enrichment data going forward could be analyses of population structure and population assignment of unknown individuals. For example, many conservation programs performing long-term biodiversity monitoring rely on reduced-SNP panels to determine provenance of an individual or sample (e.g. Bourgeois et al., 2018; Stronen et al., 2022). Developing SNP panels requires at least some *a priori* genomic resources, so target-enrichment libraries could provide feasible approaches in the absence of such references and practical improvements over existing barcoding strategies for identifying, for example, Madagascar’s rosewoods (Hassold et al., 2016); although acquiring viable DNA samples from woody tissue would remain a barrier (Jiao et al., 2018). We anticipate limited use cases of target-enrichment data in population genetics going forward as reference genome assembly is becoming more accessible through decreasing long-read costs, and less complicated library preparation protocols for whole-genome resequencing will become preferential to any target-enrichment strategy. However, the population genetic inferences from target-enrichment data should be biologically meaningful when available.

### A Practical Need for Plant Population Genomics in Madagascar

There are few studies of plant population genetics from the Central Highlands and those are limited to AFLP (Gardiner et al., 2017) or microsatellite data (Salmona et al., 2020; Hagl et al., 2021). Demographic analyses from the endemic olive tree *Noronhia lowryi* suggested recent population declines of the woody species, but it is unclear if these declines represent anthropogenic or paleoclimatic change near the LGM (Salmona et al., 2020). While our analyses of *L. simplex* are also ambiguous regarding this window of anthropogenic effects on the extent of grasslands, we can at least conclude that the *L. simplex N_e_* was quite large. Interpreting what a large *N_e_* means in the context of the *L. simplex* species distribution is difficult though; it could be a single large distribution or multiple smaller isolated distributions that yield the expected measures of genetic variation. Contemporary population genomic analyses will be necessary to provide statistical power to differentiate between competing demographic hypotheses, not only for grasses such as *L. simplex* but also species representing plant diversity across families and life-history. A better reconstruction of the natural history of Madagascar’s grasslands is not only necessary for understanding diversity within and outside of the Central Highlands (Yoder et al., 2016; Burbrink et al., 2019; Everson et al., 2020), but also to make effective land management decisions for tree planting programs.

## Acknowledgements

This project has received funding from the European Union’s Horizon 2020 research and innovation programme under the Marie Sklodowska-Curie grant agreement No 101026923, awarded to GPT and MSV. GB is supported by LabEx TULIP (ANR-10-LABX-0041). CERL was supported by a The Royal Society International Collaboration Award. Thank you to MEDD Madagascar, Madagascar National Parks, Isalo MNP, and Itremo NPA for supporting our research and exportation permits in 2014 and 2017, and also to COSTECH Tanzania for their support for the 2014 collection permits for MSV and WRQL. Sequence data were generated with research funds from Duke University to ADY. This work would not have been possible without Stuart Cable and everyone at the Kew Madagascar Conservation Centre. We thank Emily Lockwood for DNA extractions, and Lorena Endara and J. Gordon Burleigh for coordinating library preparation and sequencing. Colleen Seymour provided helpful feedback on an earlier version of this manuscript. We thank two anonymous reviewers and the editor Andrew Young for helpful comments that improved the manuscript.

## Author Contributions

GPT, ADY, and MSV conceived the study. WRQL, CLS, CERL, GB and MSV collected samples. GPT and TOMA curated data. GPT and AAC analyzed data. GPT wrote the first draft. TOMA translated the abstract to French and Malagasy. All authors revised the manuscript and approved the final version.

## Data Availability Statement

Scripts for analyses and input files are available through the Dryad Digital Repository: https://doi.org/10.5061/dryad.905qfttqx. Additional scripts are available on GPT’s GitHub for identifying exon-intron boundaries from target enrichment data (https://github.com/gtiley/Locus-Splitter) generating matrices for PCAs from fasta files (https://github.com/gtiley/fasta-conversion) and the ploidy test implemented through PATÉ (https://github.com/gtiley/Phasing). Raw sequence data is available via the NCBI short read archive (SRA) database, with individual SRA and BioSample identifiers in Supplementary Table S1. Raw sequence data is associated with NCBI BioProject PRJNA952819.

## Conflict of Interest Statement

The authors declare no conflict of interest.

## Notes

### Competing Interest Statement

The authors have declared no competing interest.

### Summary of Updates

Revisions were carried out in accordance with reviewer requests that largely concerned small text edits and clarifications. All analyses and figures remained the same. Two notable additions include: 1) Alternative language abstracts (Summary and Societal Impact Statements) in French and Malagasy. 2) Additional methods describing the identification of overlapping loci in the target-enrichment data caused by probe regions in multiple exons of the same locus. This could cause some sites to be represented multiple times in the data and potential violations of independence among loci. These biases were accounted for in the previous version, but the process for identifying and removing the overlapping loci was omitted.

