## Supplementary Information for "Genetic variation in *Loudetia simplex* supports the presence of ancient grasslands in Madagascar"

### Supplementary Methods

*Ploidy Test*

We explored the utility of normal mixture models on distributions of allele balance from variants passing filters for estimating ploidy from target-enrichment data. Allele balance is the proportion of reads with an alternate allele given the total number of observed reads for a site. The reference allele is determined by the reference genome, or in our case a reference fasta of supercontig assemblies, and the alternate allele will be an observed nucleotide change. Thus, our test only considers biallelic sites and furthermore is limited to octaploids. This is because Weiß et al. (2018) suggest it will be statistically difficult to accurate infer higher ploidy levels, and we agree.

In brief, a normal mixture model assumes there are $k$ normal (Gaussian) distributions. Each component $i \in \{1,2,\ldots,k\}$ has a mean $\mu_i$ and standard deviation $\sigma_i$ and explains a proportion $p_i$ of the total distribution. We assume that allele balance distributions can be described by normal mixtures for the following ploidy levels:

1) Diploid – A single distribution ($k$ = 1) with $p_1$ = 1 and $\mu_1$= 0.5
2) Triploid – Two distributions ($k$ = 2) with $p_1$ = 0.5 and $p_2$ = 0.5. The two distributions have means $\mu_1$= 0.33 and $\mu_2$= 0.66
3) Tetraploid – Three distributions ($k$ = 3) with $p_1$ = 0.25, $p_2$ = 0.5, and $p_3$ = 0.25. The three distributions have means $\mu_1$= 0.25, $\mu_2$= 0.5, and $\mu_3$= 0.75. Note that the proportions are unbalanced with $p_2$ = 0.5. The proportions are derived for a sexually reproducing population.
4) Pentaploid – Four distributions ($k$ = 4) with $p_1$ = 0.25, $p_2$ = 0.25, $p_3$ = 0.25, and $p_4$ = 0.25. The four distributions have means $\mu_1$= 0.2, $\mu_2$= 0.4, $\mu_3$= 0.6, and $\mu_4$= 0.8.
5) Hexaploid – Five distributions ($k$ = 5) with $p_1$ = 1/6, $p_2$ = 1/6, $p_3$ = 2/6, $p_4$ = 1/6, and $p_5$ = 1/6. The five distributions have means $\mu_1$= 1/6, $\mu_2$= 2/6, $\mu_3$= 3/6, $\mu_4$= 4/6, and $\mu_5$= 5/6.
6) Septaploid – Six distributions ($k$ = 6) with $p_1$ = 1/6, $p_2$ = 1/6, $p_3$ = 1/6, $p_4$ = 1/6, $p_5$ = 1/6, and $p_6$ = 1/6. The six distributions have means $\mu_1$= 1/7, $\mu_2$= 2/7, $\mu_3$= 3/7, $\mu_4$= 4/7, $\mu_5$= 5/7, and $\mu_6$= 6/7.
7) Octaploid – Seven distributions ($k$ = 7) with $p_1$ = 1/8, $p_2$ = 1/8, $p_3$ = 1/8, $p_4$ = 2/8, $p_5$ = 1/8, $p_6$ = 1/8, and $p_7$ = 1/8. The seven distributions have means $\mu_1$= 1/8, $\mu_2$= 2/8, $\mu_3$= 3/8, $\mu_4$= 4/8, $\mu_5$= 5/8, $\mu_6$= 6/8, and $\mu_7$= 7/8.

To potentially avoid overfitting of mixture models, all $p_i$ and $\mu_i$ were fixed to the above values for each $k_i$. The standard deviation terms of each component and the likeklihood was optimized via expectation maximization using the *optim* function from R. We used 100 independent optimizations with random starts for each model and retained the best likelihood and parameters. To determine the best $k$, and thus ploidy, for an allele balance distribution, the maximum likelihood estimates are used to calculate the model weights. This is because previous investigations showed comparisons of likelihood scores via $\Delta BIC$ could lead to false positives for higher $k$ values (Tiley et al. 2018). All code is publicly available and can be obtained from GPT's GitHub (https://github.com/gtiley/Ks_plots).

*BPP Prior Specification*

The HKY model of nucleotide substitution with gamma-distributed rate heterogeneity among sites was assumed for each alignment. The shape parameter for rate heterogeneity was draw from a gamma prior ($\alpha = 1, \beta = 1$) and discretized over four classes. A strict clock was assumed among lineages, but locus rates were allowed to vary. All locus rates were relative, such that the mean rate was 1. We chose a somewhat diffuse concentration parameter of 5, since target-enrichment data are coding but representative of a subset of typically conserved loci and assumed locus rates were distributed under a conditional independent and identically distributed prior.

The divergence time between *Loudetia* and *Tristachya* was assigned an inverse gamma prior with $\alpha = 3, \beta = 0.01$ and all other divergence times were described by a Dirichlet distribution with $\alpha = 0.01$. This implies a low expectation of genetic variation within our data, but the posterior should nevertheless pull away from the prior if data are informative. Population sizes followed an inverse gamma prior distribution with $\alpha = 2, \beta = 0.01$. This assumes that a $\theta = 0.01$ or an expected substitution very 100bp is reasonable. Although, we analytically integrated out population sizes rather than sample population size parameters via MCMC, since we were only interested in the topology for these analyses.

*PCA Input Data*
Principal component analyses (PCAs) with genetic data are a common tool for investigating population structure and are very convenient for their simplicity. However, population genomic software does not readily accommodate mixed ploidy data, although adegenet (Jombart and Ahmed 2011) can readily analyses polyploid data of the same ploidy level. Because our *Loudetia simplex* sample had likely tetraploids and hexaploids, we proposed strategies for compressing PCA matrices with higher ploidy levels to lower levels to make all individuals comparable regardless of ploidy.

The data for a PCA in population genetics is the count of observed alternate alleles. A diploid individual with two reference alleles for a site would be scored as a 0. If that individual had one reference allele and one alternate allele, it would receive a score of 1 for the site. Two alternate alleles, meaning the individual is homozygous for the alternate allele, would result in a score of 2. Now consider a hexaploid is in the analysis; it will have six alleles for a site. The hexaploidy that is homozygous for the alternate allele would receive a score of 6. This causes a problem, since a combined analysis of hexaploids and diploids would imply the diploid also homozygous for the alternate allele at a site has two alternate alleles and four reference alleles. We therefore wrote a script (https://github.com/gtiley/fasta-conversion) that can generate the following matrices that we found useful:

1) No compression (*.pcaraw.txt) – This is the raw data. It can be used directly if all individuals are the same ploidy, but it is also good to check the raw data to ensure nothing looks wrong.
2) Five category compression (*.pcacomp5.txt) – A homozygous reference site is scored as 0 and a homozygous alternate site is scored as 4. A site that is a balanced 50:50 ratio of reference to alternate alleles, regardless of ploidy, is scored as a 2. A site that is heterozygous be unbalanced with more reference than alternate alleles receives a 1. A site that is heterozygous be unbalanced with more alternate than reference alleles receives a 3. This is equivalent to a normal PCA matrix for a sample of all tetraploids.
3) Three category compression (*.pcacomp3.txt) – A homozygous reference site is scored as 0 and a homozygous alternate site is scored as 2. Any heterozygous site regardless of allelic dosage is scored as a 1. This is equivalent to a normal PCA matrix for a sample of all diploids.
4) Two category compression (*.pcacomp2.txt) – A homozygous reference site is scored as a 0 and the presence of any dosage of alternate alleles scores the site as a 1. This is equivalent to a normal PCA matrix for a sample of all haploids.

*Summary Statistics*
We were concerned if target-enrichment data could be used to estimate basic population genetic summary statistics, ignoring some model violations. The parameters we looked at were:

1) The expected pairwise nucleotide difference ($\pi$) – This can be estimated by $\frac{n}{n-1}\sum_{i=1}^{S_n} 2p_i\,(1-p_i)$, where *n* is the sample size in number of sequences, $S_n$ is the number of segregating sites for that sample, and $p_i$ is the frequency of the minor allele at site *i*. This is often called nucleotide diversity. We plotted per-site estimates by dividing the estimate of $\pi$ by the alignment length. This was done simply to provide some intuition about variation among loci. The downstream Tajima's *D* estimates do not use the per-bp scaling.
2) The number of segregating sites (*S*) – This is simply the number of alignment columns that have a non-trivial difference (not a gap or missing data). Population geneticists are interested in *S* because it is a sufficient statistic for Watterson's estimator of nucleotide diversity ($\theta$; Watterson 1975). Watterson's estimator is $\theta = \frac{S_n}{\sum_{i=1}^{n-1}\frac{1}{i}}$ , such that $S_n$ is the number of segregating sites in a sample of *n* sequences. *S* is reported for each alignment without any rescaling.
3) Deviation between unbiased estimators of nucleotide diversity (Tajima's *D*) – Both $\pi$ and $\theta$ are unbiased estimators of nucleotide diversity and a population that meets assumptions of neutrality and constant population size, among others, should result in the same values between the two estimators. Differences between the two estimators reveals deviations from neutrality. Tajima (1989) analytically solved a meaty normalizing constant not shown here to leads to interpretations of Tajima's *D* less than -2 as more rare alleles than expected are in the population (possibly due to strong directional selection or population size increase). Tajima's *D* greater than 2 implies fewer rare variants are in the population than expected (possibly due to balancing selection or population size decrease). Because we do not have alignments of infinite length and there is stochasticity in the coalescent process dependent on the number of individuals, we used simulations to generate *p*-values for each alignment, to ensure any one Tajima's *D* statistic would not be due to chance alone.

   The ms commands used for simulating the null distribution of Tajima's *D* for the different data sets are:

   OH) `ms 20 1000 –t 1.14210526315789`
   AG) `ms 40 1000 –t 1.61923076923077`
   PH) `ms 120 1000 –t 0.957422969187675`

   Such that the `–t` parameter is the average locus-scaled $\theta$ calculated by the PopGenome v2.7.5 R package (Pfeifer et al., 2014).

To investigate the stability of the summary statistics, we generated two terms that are not used correctly from a statistical standpoint but are perhaps useful. First, we looked at the bias between the observed estimate and the mean bootstrapped estimate. This should provide some estimate of the uncertainty in our population summary statistics due to sampling error. A very small bias implies the sequences are long enough and that there is likely little to be gained from sampling more sites at a locus. We also explored the root squared mean error (RSME) calculated by $\sqrt{\frac{\sum_{i=1}^{n}(x_i-x)^2}{n}}$ where *x* is the observed estimate and $x_i$ is a bootstrap estimate for *n* bootstrap replicates. This should provide some intuition on the variation of estimates among bootstrap replicates and if there are differences in uncertainty based on how ploidy is treated. Both terms were reported in lieu of bootstrap confidence intervals around the observed estimate, because the observation typically fell outside of the confidence interval. This is likely due to the skewed nature of distributions of genetic variation, assuming mutations arise from a Poisson process.

*EBSP Prior Specification*
The HKY model of nucleotide substitution with gamma-distributed rate heterogeneity among sites was assumed for each alignment. The transition-transversion rate ratio was drawn from a lognormal prior with mean of 1 and standard deviation of 1.25. The shape parameter for rate heterogeneity was draw from a gamma prior ($\alpha = 1, \beta = 1$) and discretized over four classes. Empirical equilibrium frequencies were used. All tree priors used the coalescent extended Bayesian skyline (Heled and Drummond 2008). In an attempt to provide some numerical stability to the MCMC search, we set the clock rate to 0.001, although we anticipate a per-year substitution rate of $1 \times 10^{-9}$ (Christin et al. 2014). This scaling resulted in reproducibility among identical runs, within reason of MCMC proposals. Therefore, the effective population size parameter was drawn from an exponential distribution with a mean of 3, which is reasonably vague while allowing exploration of some higher values.

There are may unknowns in the life history of *Loudetia simplex*. For example, there may be clonal propagation from rhizomes, asexual reproductions, and certainly overlapping generations among sexually reproductive individuals. Thus, we do not attempt to characterize a generation time for the species. We assume $1 \times 10^{-9}$ is a reasonable starting point for perennial panicoid bunch grasses, but introduce uncertainty through a rate prior. We assume the rate is gamma distributed with $\alpha = 10$ and $\beta = 10000$. This centers the distribution on 0.0001, but nearly allows for a two-order-of-magnitude uncertainty about the rate. Although heavily centered on our initial rate assumption, we can explore the effects of rate uncertainty on the ultimate biological interpretations of analyses and draw some conclusions about how wrong we would have to be about the rate to be wrong about the conclusions.

Similar to our BPP analyses, we allowed rate variation among loci. All locus rates were relative to a single locus, which was calibrated to the clock rate. This was a choice made to place some constraints on the model and ensure that rates were statistically identifiable. Although this places a lot of weight on the reference locus calibrated to the clock rate, exploration of a few loci revealed that analyses were not very sensitive to the reference locus, at least for our data. XML files for reproducing analyses and observing model specifications are available on Dryad (X).

*Simulating Demographic Histories*
Simulations, while not capturing the complexities of real data, can be a useful tool to evaluate the limits of inference methods under best-case scenarios. We are skeptical that the EBSP or related methods could reliably differentiate anthropogenetic- versus paleoclimate-associate changes in effective population size, especially in a large population organism like grasses. We explored some simple scenarios with Hudson's ms (Hudson 2002) and provide some details about the simulation commands here. All simulations were for 20 individuals (haplotype sequences), 57 loci (our empirical sample),1000 bp loci (close enough to our average alignment length), and no recombination.

1. A constant population size

A single diploid population with $N_e$ of 100,000 was simulated under the neutral coalescent. The -t parameter of ms is per-locus scaled nucleotide diversity, such that $\theta_L = 4N_e\mu_L$ where $\mu_L$ is the per locus mutation rate as opposed to a per-bp mutation rate. Here $\theta_L = 4 \times 100{,}000\ individuals\ \times (1 \times 10^{-8}\ mutations\ per\ generation\ \times 1{,}000\ bp) = 4$. Then, *i* simulations from 1 to 100 where the seeds are provided by Perl's default *rand* function looks like:

```
ms 20 57 -t 4.0 -T -seeds $seed1 $seed2 $seed3 -r 0 1000 | grep -F
\";\" > $i.20.treefile
seq-gen -mHKY -l 1000 -s 0.004 -p 1 -a 0.7 -f 0.3,0.2,0.2,0.3 -t 1.5 <
$i.20.treefile > $i.20.phylip
```

2. An LGM expansion

Effective population size increased from 10,000 to 100,000 individuals at 20,000 years in the past. Assuming a generation time of 10 years, this means the increase happened 2,000 generation in the past. When the time in generations is scaled by the contemporary population size, this gives the event time for ms at 0.005, when the current $N_e$ was resized to the past $N_e$ by a factor of 0.1

```
ms 20 57 -t 4.0 -eN 0.005 0.1 -T -seeds $seed1 $seed2 $seed3 -r 0 1000
| grep -F \";\" > $i.20.treefile
seq-gen -mHKY -l 1000 -s 0.004 -p 1 -a 0.7 -f 0.3,0.2,0.2,0.3 -t 1.5 <
$i.20.treefile > $i.20.phylip
```

3. An Anthropocene expansion

Effective population size increased from 10,000 to 100,000 individuals at 2,000 years in the past. Assuming a generation time of 10 years, this means the increase happened 200 generation in the past. When the time in generations is scaled by the contemporary population size, this gives the event time for ms at 0.0005, when the current $N_e$ was resized to the past $N_e$ by a factor of 0.1

```
ms 20 57 -t 4.0 -eN 0.0005 0.1 -T -seeds $seed1 $seed2 $seed3 -r 0
1000 | grep -F \";\" > $i.20.treefile
seq-gen -mHKY -l 1000 -s 0.004 -p 1 -a 0.7 -f 0.3,0.2,0.2,0.3 -t 1.5 <
$i.20.treefile > $i.20.phylip
```

4. Continuous growth from the LGM to present

The true $N_e$ history can be complex, a compound process of paleoclimate, anthropogenetic effects, and unaccounted sources. This required an additional exponential growth rate parameter for ms. Consider the basic exponential growth function where $N_1 = N_0 e^{-\alpha t}$ and $N_1$ is the contemporary size of 100,000 and $N_0$ is the past size of 10,000. Our event time $t$ is already provided for 20,000 years or 2,000 generations ago, 0.005. This gives $\alpha = -\frac{log\left(\frac{N_1}{N_0}\right)}{t} = -\frac{\log(10)}{0.005} = -460.517$.

```
ms 20 57 -t 4.0 -eN 0.005 0.1 -G 460.517 -T -seeds $seed1 $seed2
$seed3 -r 0 1000 | grep -F \";\" > $i.20.treefile
seq-gen -mHKY -l 1000 -s 0.004 -p 1 -a 0.7 -f 0.3,0.2,0.2,0.3 -t 1.5 <
$i.20.treefile > $i.20.phylip
```

### Supplementary Figures

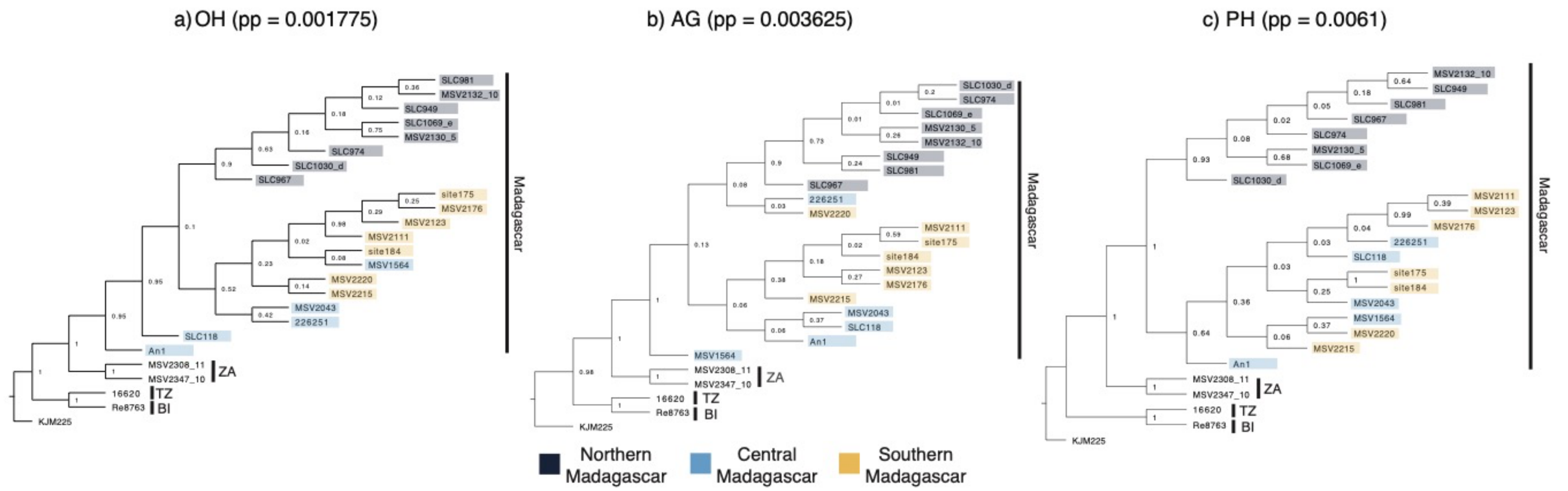


**Supplementary Figure S1 — Species trees from BPP.** The posterior probability (pp) for the tree is given for each data type based on 40,000 posterior samples. Node values are posterior probabilities for bipartitions in the posterior sample. Geographic regions are colored subjectively based on STRUCTURE results and sampling site.

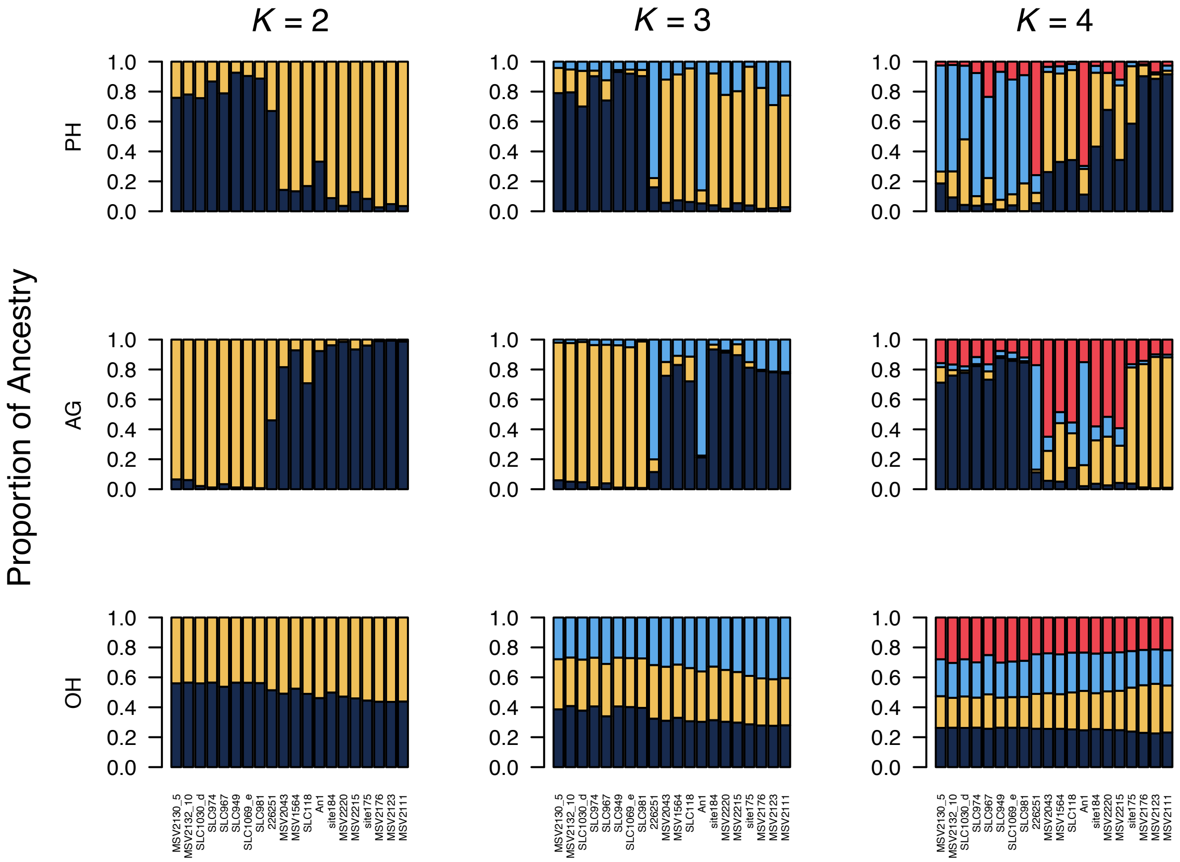


**Supplementary Figure S2 — STRUCTURE results with increasing *K*.** Cluster assignments are weighted across biological and technical replicates.

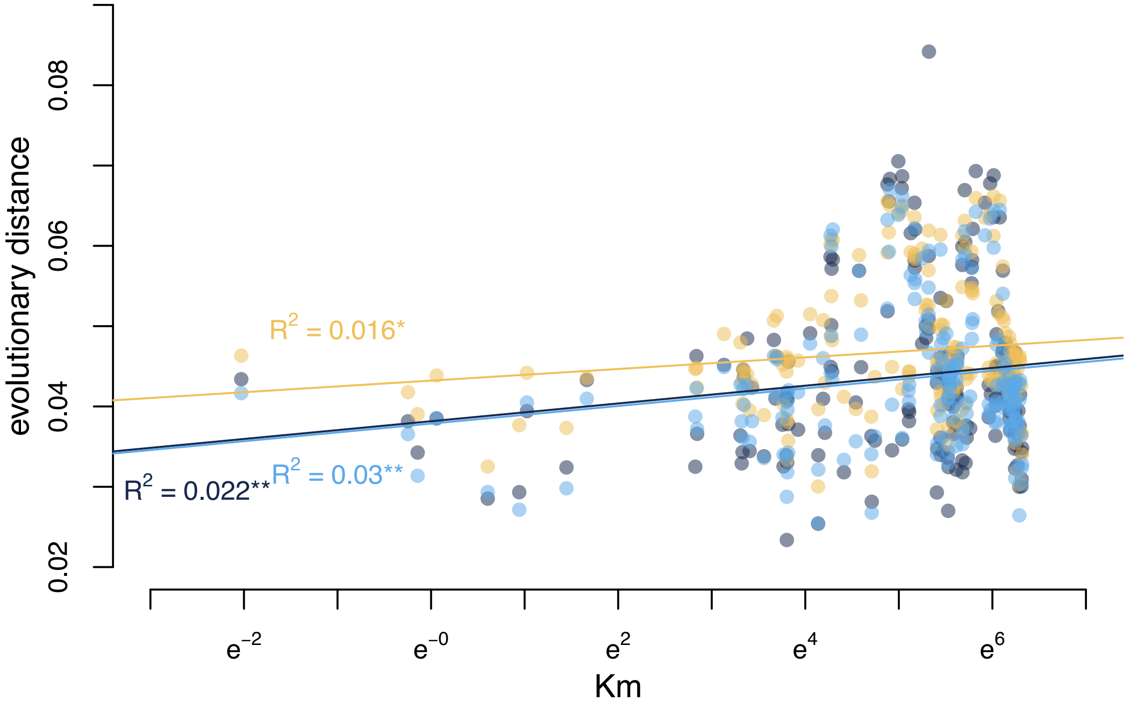


**Supplementary Figure S3 — Isolation-by-distance with evolutionary distances from ML phylogenetic analyses.** A Mantel test was used to assess statistical significance. p<0.05*, p<0.01**, p<0.001***.

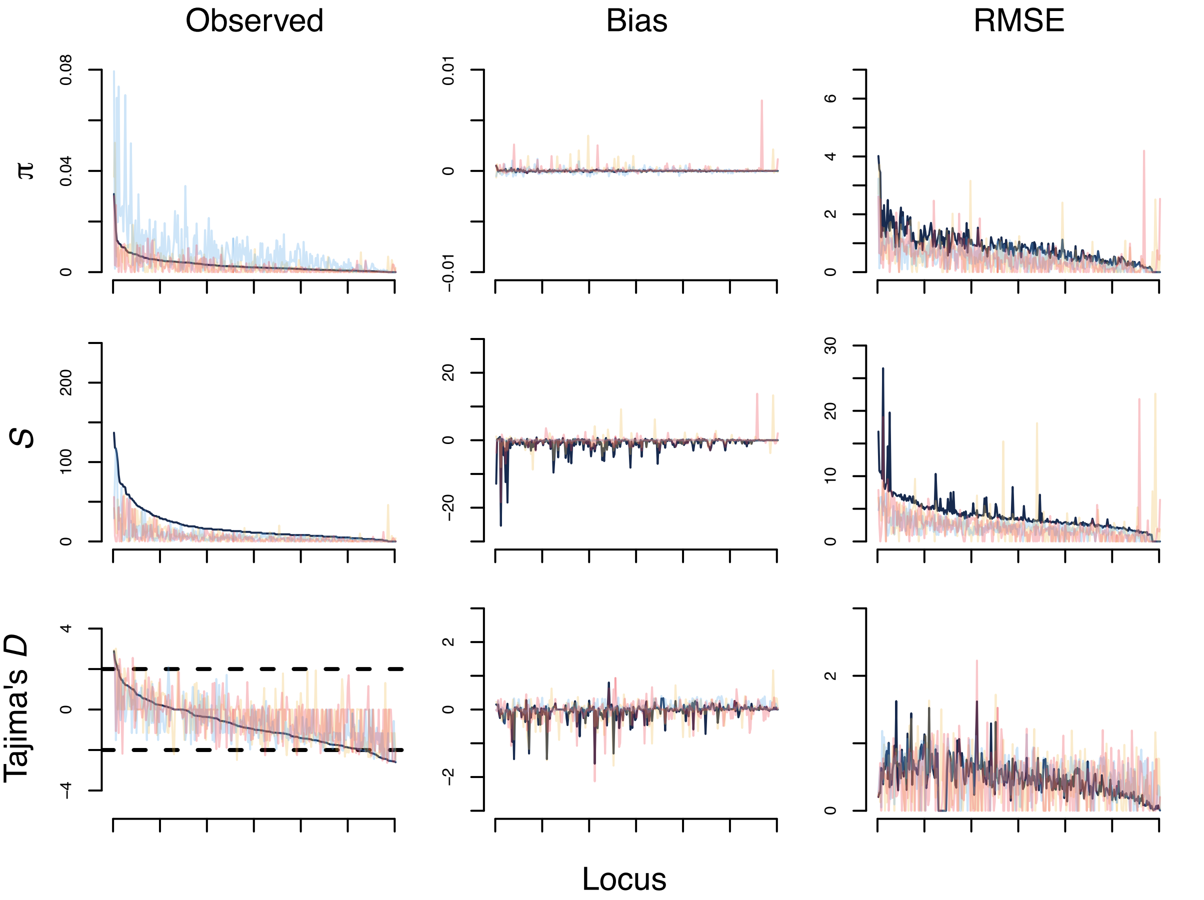


**Supplementary Figure S4 — Distributions of summary statistics across OH loci.** Loci are ordered by value from the whole-locus estimates.

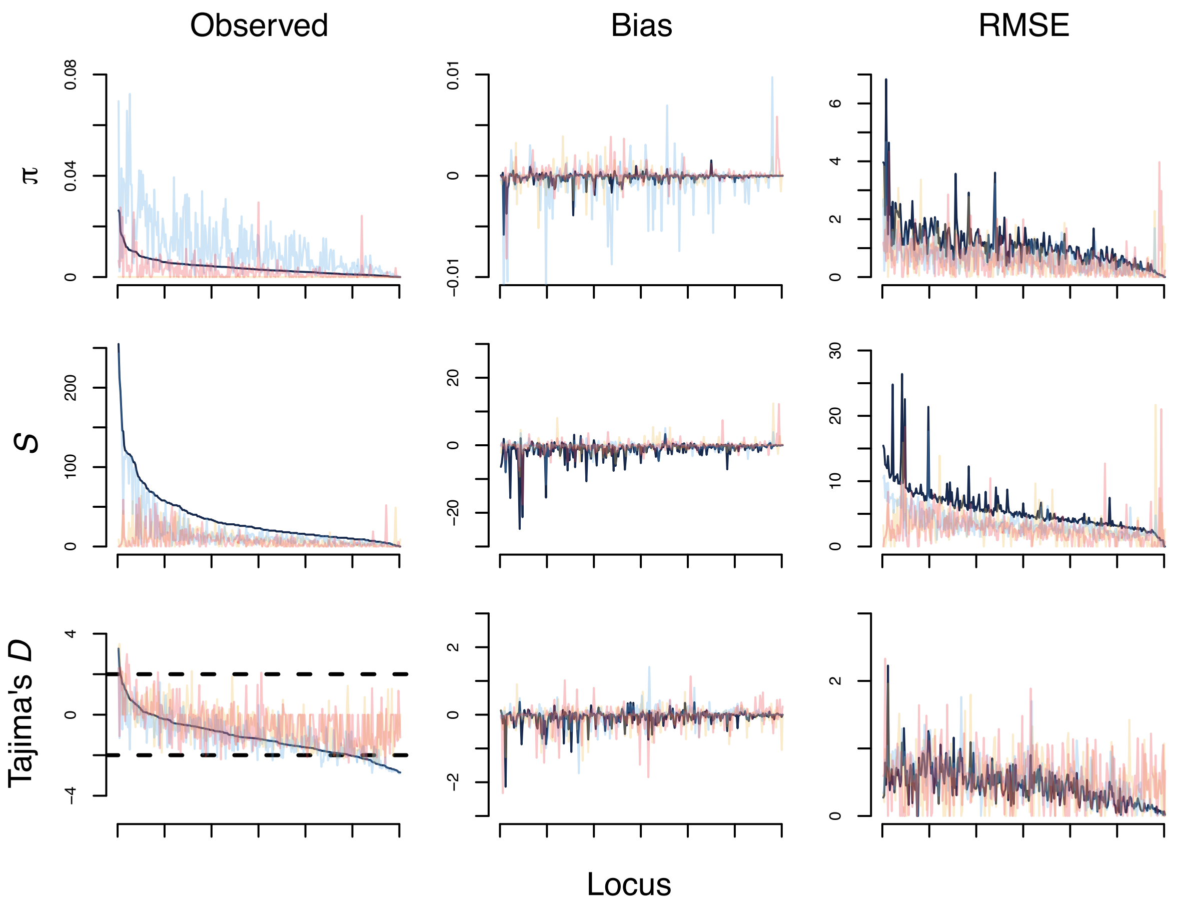


**Supplementary Figure S5 — Distributions of summary statistics across AG loci.** Loci are ordered by value from the whole-locus estimates.

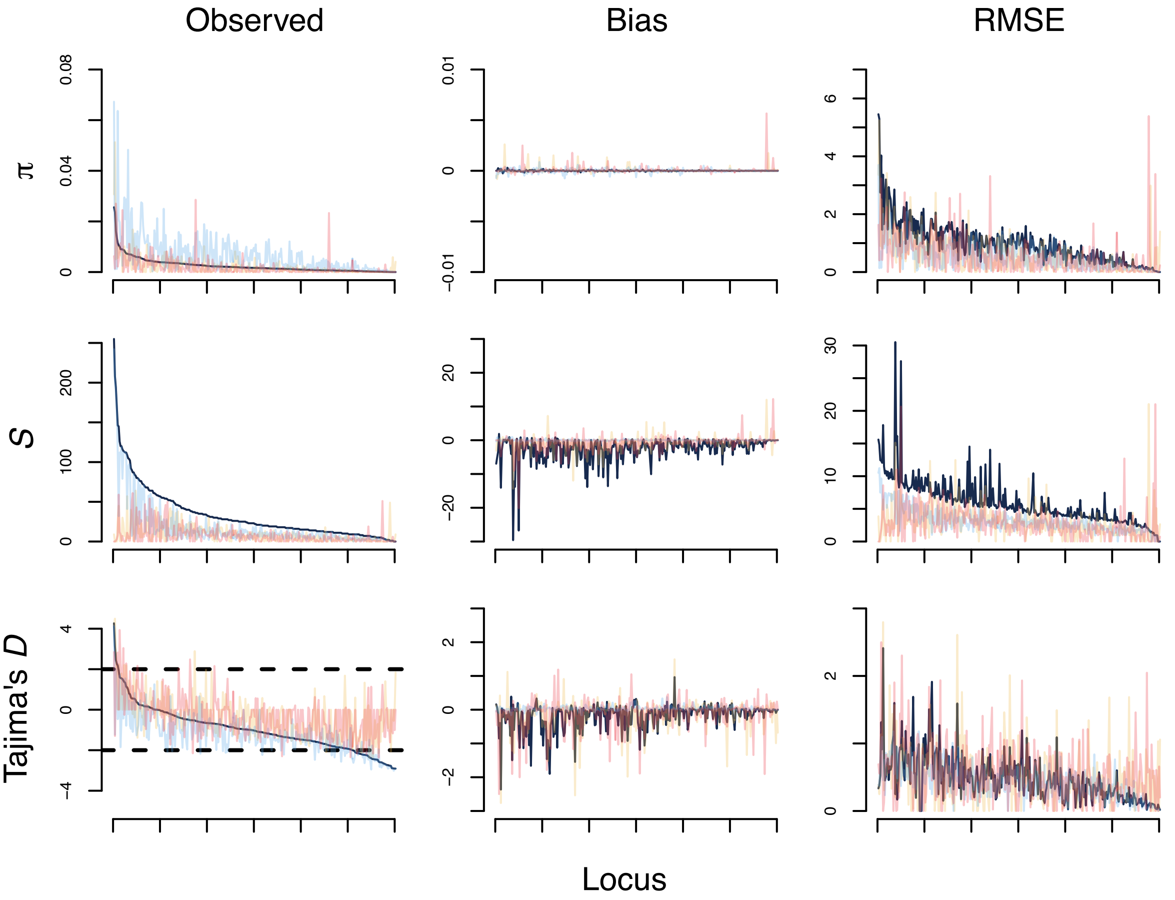


**Supplementary Figure S6 — Distributions of summary statistics across PH loci.** Loci are ordered by value from the whole-locus estimates.

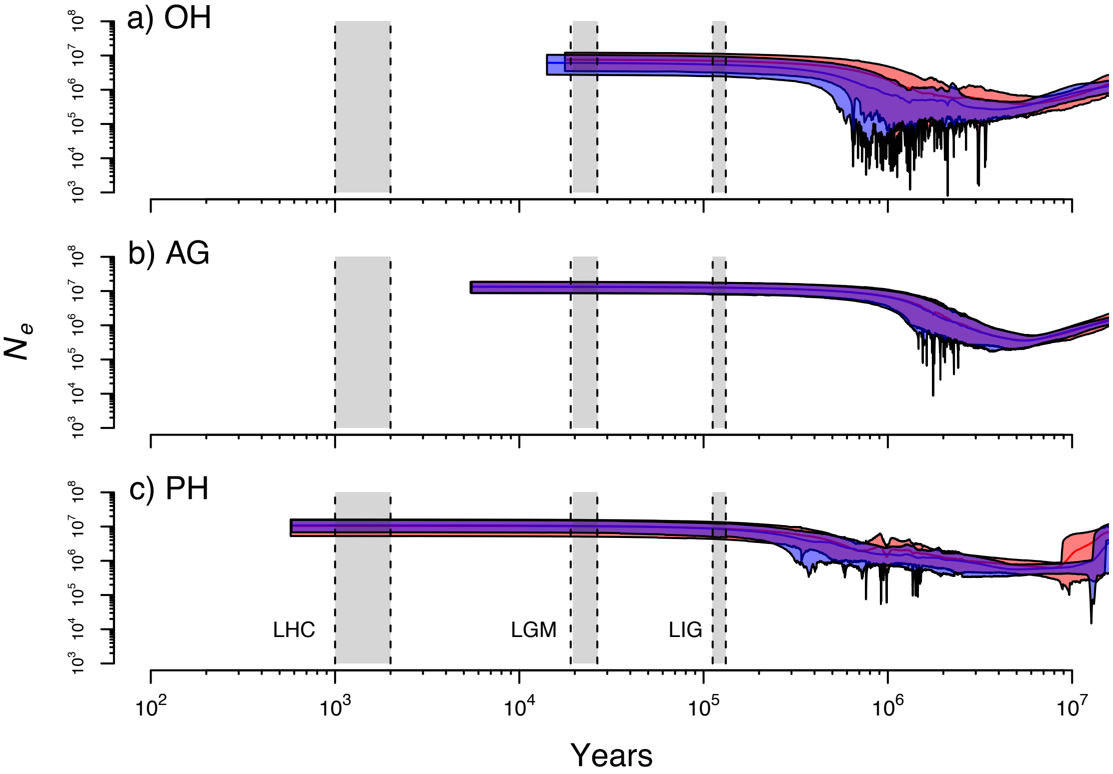


**Supplementary Figure S7 — EBSP results with variable clock rates.** Two independent runs are shown with mean population sizes shown by solid lines. Polygons show the 95% HPD intervals with the overlap between runs in purple. Results are shown for the a) OH, b) AG, and c) PH data. Relevant time periods are shown in gray, which includes likely human colonization (LHC), the last glacial maximum (LGM), and last interglacial period (LIG).

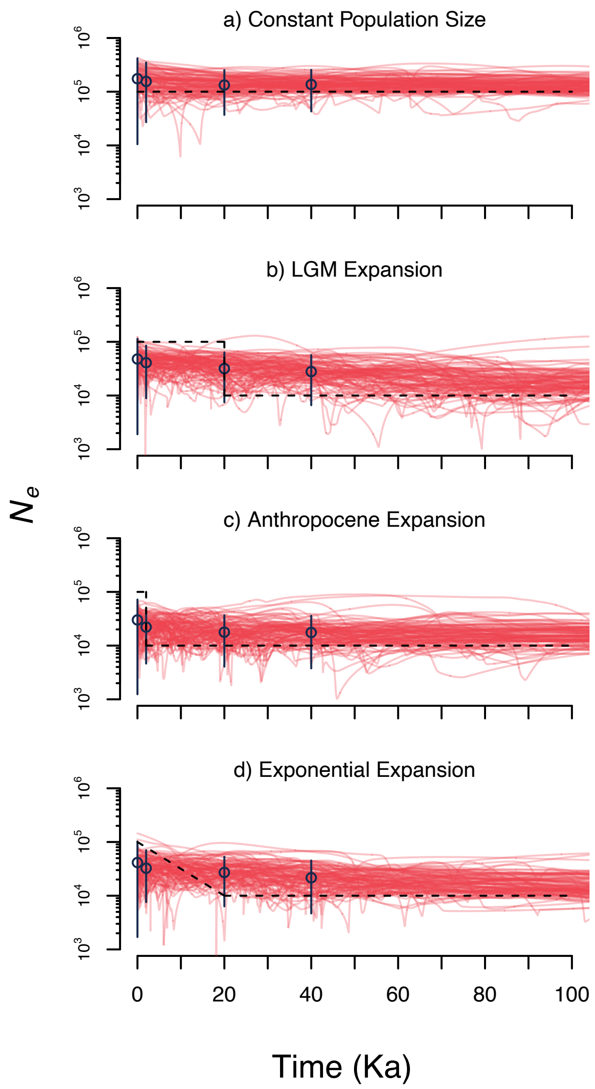


**Supplementary Figure S8 — Simulation results from EBSP analyses for a misspecified high number of change points.** Individual red lines are the mean $N_e$ estimates from 100 simulated data sets. Blue points and vertical lines are the mean of mean $N_e$ estimates and their 95% HPD intervals. The dashed black line is the true demographic history of the population.

### Supplementary Tables

Supplementary tables are provided in a separate excel spreadsheet for ease of use.

**Table S1 — Metadata for sampled individuals.** The anticipated ploidy is based on the maximum number of alleles observed across nuclear microsatellite loci from Hagl et al. (2021).

**Table S2 — Sequencing and phasing summary statistics from target-enrichment loci.** A block is a contiguous phased sequence. Phasing and read statistics were calculated before removing loci suspected of being in tight linkage.

**Table S3 — Mixture model results from allele balance distributions.** The estimated ploidy is the number of inferred components + 1. The mixing proportions (p) and means (m) are fixed for each component while the standard deviation (sd) is freely estimated.

**Table S4 — STRUCTURE results across 20 runs for one to five clusters (K) for 10 data matrices that randomly sampled 1 SNP per locus.** The delta K estimate and intermediate calculations are shown for each analysis of 20 runs.

**Table S5 — Differences in distributions of summary statistics between data type and gene region.** Whole-locus comparisons were made between the OH, AG, and PH data, but not to sub-regions. P-values from Wilcoxon rank-sum tests are on the upper diagonal and significance after a Bonferroni correction is given on the lower diagonal. Non-significant (ns), p<0.05 (*), p<0.01 (**), p<0.001(***).

**Table S6 — Summary statistics across locus region (including the whole locus) and data type.** The mean, percentile bootstrap 95% CIs, RMSE, and Bias are based on 200 bootstrap replicates. The probability for individual Tajima's D values are based on 100 simulations under the standard neutral coalescent model.
